# The proteomic landscape of synaptic diversity across brain regions and cell types

**DOI:** 10.1101/2023.01.27.525780

**Authors:** Marc van Oostrum, Thomas Blok, Stefano L. Giandomenico, Susanne tom Dieck, Georgi Tushev, Nicole Fürst, Julian Langer, Erin M. Schuman

## Abstract

Brain function relies on communication via neuronal synapses. Neurons build and diversify synaptic contacts using different protein combinations that define the specificity, function and plasticity potential of synapses. More than a thousand proteins have been globally identified in both pre- and postsynaptic compartments, providing substantial potential for synaptic diversity. While there is ample evidence of diverse synaptic structures, states or functional properties, the diversity of the underlying individual synaptic proteomes remains largely unexplored. Here we used 7 different Cre-driver mouse lines crossed with a floxed mouse line in which the presynaptic terminals were fluorescently labeled (SypTOM) to identify the proteomes that underlie synaptic diversity. We combined microdissection of 5 different brain regions with fluorescent-activated synaptosome sorting to isolate and analyze using quantitative mass spectrometry 18 types of synapses and their underlying synaptic proteomes. We discovered ~1’800 unique synapse type-enriched proteins and allocated thousands of proteins to different types of synapses. We identify commonly shared synaptic protein modules and highlight the hotspots for proteome specialization. A protein-protein correlation network classifies proteins into modules and their association with synaptic traits reveals synaptic protein communities that correlate with either neurotransmitter glutamate or GABA. Finally, we reveal specializations and commonalities of the striatal dopaminergic proteome and outline the proteome diversity of synapses formed by parvalbumin, somatostatin and vasoactive intestinal peptide-expressing cortical interneuron subtypes, highlighting proteome signatures that relate to their functional properties. This study opens the door for molecular systems-biology analysis of synapses and provides a framework to integrate proteomic information for synapse subtypes of interest with cellular or circuit-level experiments.

## Introduction

The compartmentalization of biological processes in space and time within the single cell enables parallel processing of many reactions. Neurons, highly polarized cells, represent an extreme example of parallel processing – at their synapses they compartmentalize information processing to communicate with thousands of other neurons (Hanus and Schuman 2013). Synapses are plastic in structure and function, depending on their developmental and experiential history. This plasticity provides a molecular basis for learning and memory formation (Magee and Grienberger 2020).

The identity, copy number and interactions of individual proteins largely determine the physiological properties of a given synapse. Alternative splicing and post translational modifications further increase the molecular complexity of synapses. As a result, there is considerable potential for synapse molecular diversity originating from variance in proteome composition and the associated protein interaction networks. A growing body of evidence has found substantial structural and functional diversity of synapses (O’Rourke et al. 2012), but the underlying diversity in synaptic molecular architecture is much less understood. As synaptic contacts are fundamental for development and plasticity of neuronal circuits, a detailed understanding of synapse molecular diversity allows us to link molecular architecture to functional traits and disease phenotypes (Nusser 2018). Indeed, all neurodegenerative and neuropsychiatric disorders are associated with alterations in synaptic proteins (Grant 2012, 2019), and a detailed understanding of the molecular underpinnings of diversity in synapse organization will open up new rational avenues for therapeutic intervention.

Synapses transmit information by the presynaptic release of neurotransmitters, which diffuse and bind to receptors anchored in the membrane of their postsynaptic partners. This process requires a complex molecular machinery including proteins that associate with synaptic vesicles and regulate the release process, proteins that serve as scaffolds or regulators of receptors, and a diverse assortment of regulatory enzymes. Altogether, synaptic proteins work together to integrate, adjust and fine-tune the activity of the synapse to different internal and external stimuli. The complement of proteins that comprise average synaptic and sub-synaptic structures, such as the postsynaptic density or synaptic vesicles, have been studied, but synaptic proteome diversity with regards to different types and states remains largely unexplored (Koopmans et al. 2019), mainly due to limitations in technologies to analyze the proteomes of different synapse populations.

Imaging techniques provide the spatial resolution required to distinguish individual synapses and have illustrated substantial synaptic diversity for selected proteins (Zhu et al. 2018; Cizeron et al. 2020; S.-M. Guo et al. 2019; Upmanyu et al. 2022), but their limited multiplexing capabilities preclude the study of synaptic proteomes. The traditional biochemical isolation of synaptic terminals or synaptic elements (Takamori et al. 2006; Bayés et al. 2011; Roy et al. 2018; Wang et al. 2022; Wilhelm et al. 2014) yields synapse-enriched fractions that can be analyzed by mass spectrometry-based proteomics. These preparations, however, are often of low purity and contain a heterogenous mixture of many synapse types along with non-synaptic contaminants. Immunoisolation and proximity-labeling strategies coupled to mass spectrometry have been used to identify proteins residing at selected synaptic sub-compartments of defined synapse types (Boyken et al. 2013; Loh et al. 2016; Uezu et al. 2016; Spence et al. 2019). In order to isolate a defined synapse population containing all synaptic elements from *in vivo* brain structures, Biesemann et. al. introduced fluorescence-activated synaptosome sorting (FASS) (Luquet et al. 2017; Paget-Blanc et al. 2022; Hobson, Kong, et al. 2022; Hafner et al. 2019; Biesemann et al. 2014). This strategy utilizes fluorescent labeling of a synaptic protein to sort synaptosomes with high purity using a fluorescence-activated cell sorter and has been used to identify proteins that are enriched at growth cones (Poulopoulos et al. 2019) or selected synapse types (Paget-Blanc et al. 2022; Biesemann et al. 2014; Apóstolo et al. 2020).

Here, we used Cre-inducible knock-in mice and FASS coupled to mass spectrometry to systematically investigate the diversity of the synaptic proteome across genetically-defined synapse types and brain areas. We thereby outline a roadmap for molecular systems-biology analysis of synapses and identify >1’800 unique synapse-enriched proteins that compose the 18 synapse-type specific proteomes. We obtain a brain-wide protein-protein correlation network that reveals excitatory and inhibitory synaptic protein communities and highlights core and auxiliary proteins associated with the neurotransmitter glutamate and GABA. This resource reveals commonly shared synaptic protein modules as well as proteins that customize the proteomes of synapse populations in relation to their function, for example at dopaminergic synapses in the striatum and subtypes of cortical interneurons.

## Results

We developed a streamlined workflow to quantify the proteomes of synapses formed by different cell types in different brain areas. We used various Cre-inducible knock-in mice expressing a fluorescently-labeled presynaptic protein, synaptophysin-TdTomato, (SypTOM; Ai34D, JAX no: 012570) to target synapses that arise from cell types that use different neurotransmitters (e.g. excitatory, inhibitory, modulatory). We prepared and purified synaptosomes using FASS and coupled the output to a mass spectrometer to reproducibly quantify thousands of proteins **(Figure 1A)**. We first tested the pipeline by comparing well-characterized Gad2-(cortical inhibitory) and Camk2a-cre (cortical excitatory) Cre-driver lines (Taniguchi et al. 2011; Tsien, Huerta, and Tonegawa 1996) crossed with SypTOM mice **(Figure 1A)**. We prepared synaptosomes from cerebral cortex using an established Percoll-gradient procedure (Westmark et al. 2011; Dunkley, Jarvie, and Robinson 2008) and verified that synaptosome fractions were enriched in pre- and postsynaptic proteins and depleted for non-synaptic contaminants **(Figure S1).** We optimized the FASS (Biesemann et al. 2014; Hafner et al. 2019; Luquet et al. 2017; Paget-Blanc et al. 2022) strategy to define a population (P3) with a high TdTomato signal in combination with high membrane content, visualized by an FM dye **(see Methods)**. While control synaptosomes prepared from wild-type mice showed no detectable particles in the P3 gate, Camk2a∷SypTOM synaptosomes constituted approximately 40% of the total particles **(Figure 1B)**. Re-analysis of sorted synaptosomes by flow cytometry routinely led to a purity of ~80-95% **(Figure 1B)**. Immunolabeling of individually spotted sorted synaptosomes revealed that the majority of synaptosomes contain a postsynaptic element, ~65% of Camk2a∷SypTOM synaptosomes possessed labeling for the excitatory postsynaptic marker PSD-95 while ~50% of the Gad2∷SypTOM synaptosomes possessed labeling for the inhibitory postsynaptic marker gephyrin **(Figure S2)**. In this proof-of-principle experiment, we sorted ~20 million cortical synaptosomes from 8 Camk2a∷SypTOM and 8 Gad2∷SypTOM mice and quantified their proteins using mass spectrometry (Gillet et al. 2012; Muntel et al. 2019). For each mouse we additionally processed control samples consisting of the same number of particles that showed signal above a membrane dye threshold **(see methods)**. Overall, we quantified >2,300 protein groups. We defined the individual synaptic proteomes as the complement of proteins that were significantly enriched relative to their respective control fractions **(see methods, Table S1, Tab1 & Tab2)**. We found that the proteomes from Camk2a∷SypTOM and Gad2∷SypTOM synaptosomes were clearly separated from one another in a principal component analysis **(Figure 1D)**. Additionally, when comparing the sorted synaptosome fraction with unsorted control synaptosomes, we detected a specific enrichment of synaptic proteins and a depletion of mitochondrial and glial contaminants **(Figure 1E)**. We examined the proteins with the highest enrichment in the direct quantitative comparison of Camk2a∷SypTOM and Gad2∷SypTOM proteomes and observed that they represent almost exclusively established marker proteins for cortical excitatory and inhibitory synapses **(Figure 1E and Figure 1F, Table S1, Tab 3)**. We compared the 20 most enriched proteins (top ~10%) for Camk2a∷SypTOM with the SynGO synaptic protein database (Koopmans et al. 2019) and found 19 with prior synaptic annotation. These data demonstrate that this workflow can identify synaptic proteomes from a small number of purified synapses. The direct comparison of excitatory and inhibitory synaptosomes constitutes the first in-depth quantitative analysis of two distinct types of synaptic proteomes and the first purification and proteome analysis of inhibitory synaptosomes.

**Figure 1.**
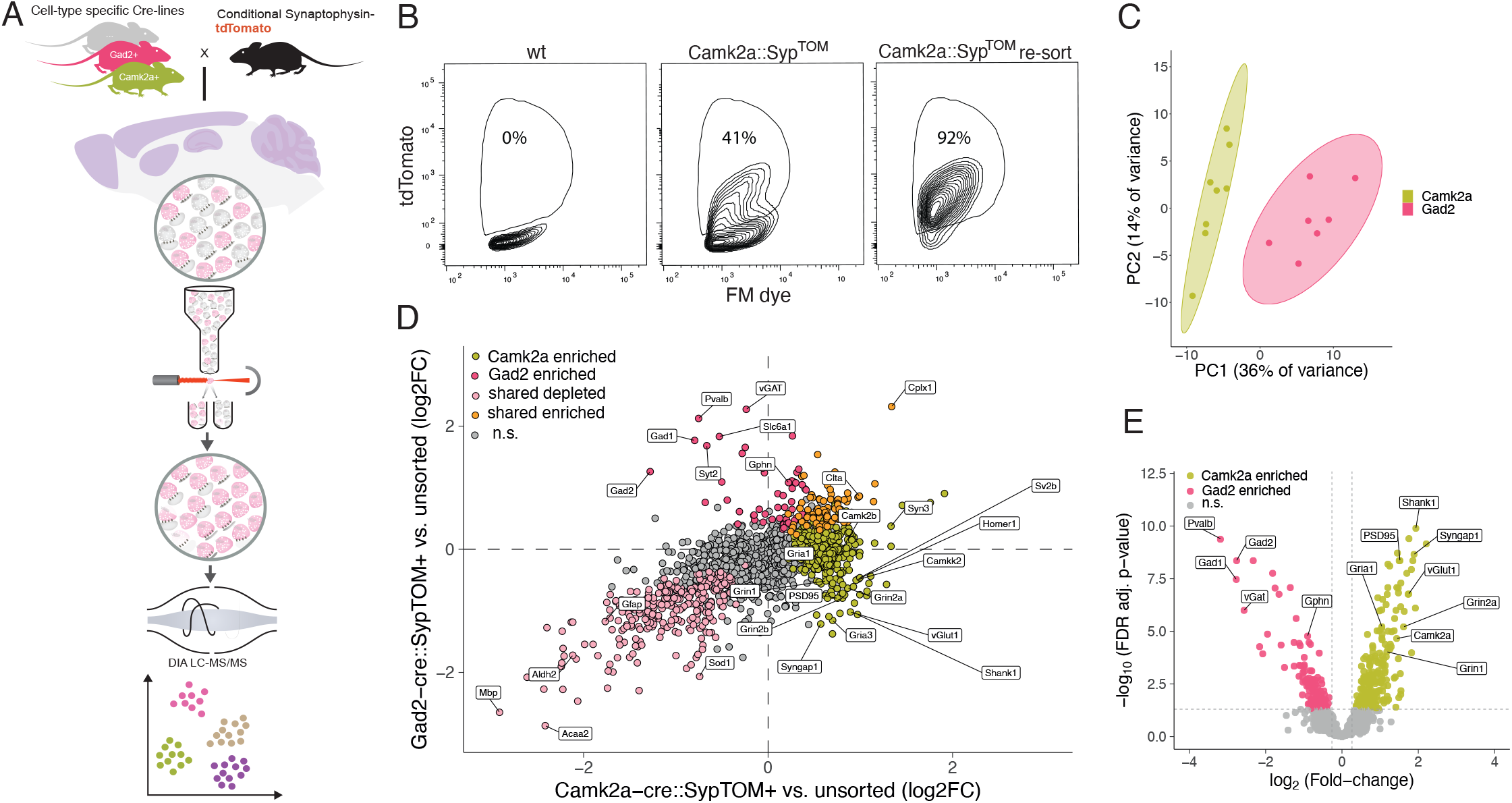
Synaptic diversity proteomic discovery pipeline and proof-of-principle. (A) The pipeline begins with the crosses of different cell type-specific Cre driver lines (Camk2a+, Gad2+, Syn1+, Dat+, PV+, SST+, VIP+) and a floxed Synaptophysin-TdTomato line, resulting in the cell type-specific labeling of presynaptic terminals. Different indicated brain regions (olfactory bulb, cortex, striatum, hippocampus and cerebellum) were microdissected and synaptosomes were generated from each region. Fluorescence-activated synaptosome-sorting (FASS) was used to purify the fluorescent, cell type-specific population of synaptosomes from each area. Then each purified synaptosome population was subjected to data-independent acquisition (DIA) mass spectrometry and the proteomes determined by statistical analysis of quantitative enrichment. (B) Gating strategy and sorting efficiency. FASS contour plots showing the relative density of the targeted TdTomato+ synaptic population in cortical synaptosomes prepared from wild-type mice (0%; left) or Camk2a+-Cre:SypTom mice (41%; middle). X-axis represents fluorescence from a membrane dye (see methods) and the y-axis represents fluorescence from TdTomato. Following the initial sorting run (middle), re-loading of the sorted synaptosomes indicated a high enrichment and purity (92%) of the Camk2a+-Cre:SypTOM sample (right). (C) Principal components analysis (PCA) showing the clear separation of Camk2a+ vs. Gad2+-sorted synaptosome proteomes. (D) Scatter plot comparing the differential enrichment of proteins in the Camk2a+-sorted and Gad2+-sorted synaptosomes to their control synaptosome precursor populations. Indicated are proteins that are significantly enriched in Camk2a+-sorted synaptosomes (lime green), Gad2+-sorted synaptosomes (rose), significant in both populations (orange) and significantly de-enriched in both (pale pink). Note the specific enrichment of labeled marker proteins for excitatory and inhibitory proteins. (E) Differential enrichment Volcano plot comparing the proteins significantly enriched in Camk2a+-sorted synaptosomes (lime green), vs. Gad2+-sorted synaptosomes (rose). Some canonical marker proteins for excitatory and inhibitory synapses are highlighted.

We applied the above pipeline to investigate the proteomic diversity of 15 different major synapse subtypes using four Cre-driver lines representing different cell types and microdissection of five different brain areas (cortex (CX), hippocampus (HC), striatum (STR), olfactory bulb (Bulb) and cerebellum (CER))**(Figure 2A)**. Besides Camk2a- and Gad2-cre, we included Syn1-cre, a line which is independent of neurotransmitter type, and Dat-cre, representing the synapses that use the modulatory neurotransmitter dopamine. In a series of control experiments, we first comprehensively assessed the degree to which a given Cre-line labeled the expected synaptic population in a given brain area. We characterized the neurons and the synapses expressing the SypTOM construct using fluorescent *in situ* hybridization (FISH) **(Figure S3)** and immunofluorescence **(Figure S4)** for canonical glutamatergic and GABA-ergic markers, and we made use of relevant single-cell sequencing data by Zeisel et. al. (Zeisel et al. 2018) **(Figure S5)**. In summary, we found that in the Cortex and Hippocampus Camk2a∷SypTOM and Gad2∷SypTOM mice largely label mutually exclusive neuron types expressing either vGlut1 or Gad2 mRNA or protein. In the Striatum and Olfactory bulb, however, Camk2a∷SypTOM and Gad2∷SypTOM label both distinct and overlapping neuron populations **(Figure S3-5)**. In Syn1∷SypTOM mice we detected TdTomato mRNA in both vGlut1- and Gad2-expressing neurons, but, as observed by others (Paton et al. 2022), often only a subpopulation is labeled as compared with Camk2a∷SypTOM or Gad2∷SypTOM. Finally, we examined by 2D electron microscopy whether synaptosomes from different brain regions differ in their ultrastructural features. We found no significant differences in synaptosome abundance, in the size or presence of synaptic vesicles, or in synaptic mitochondria or postsynaptic structures between brain regions **(Figure S6)** and we verified that synaptosome ultrastructure remains intact after sorting **(Figure S7)**.

**Figure 2.**
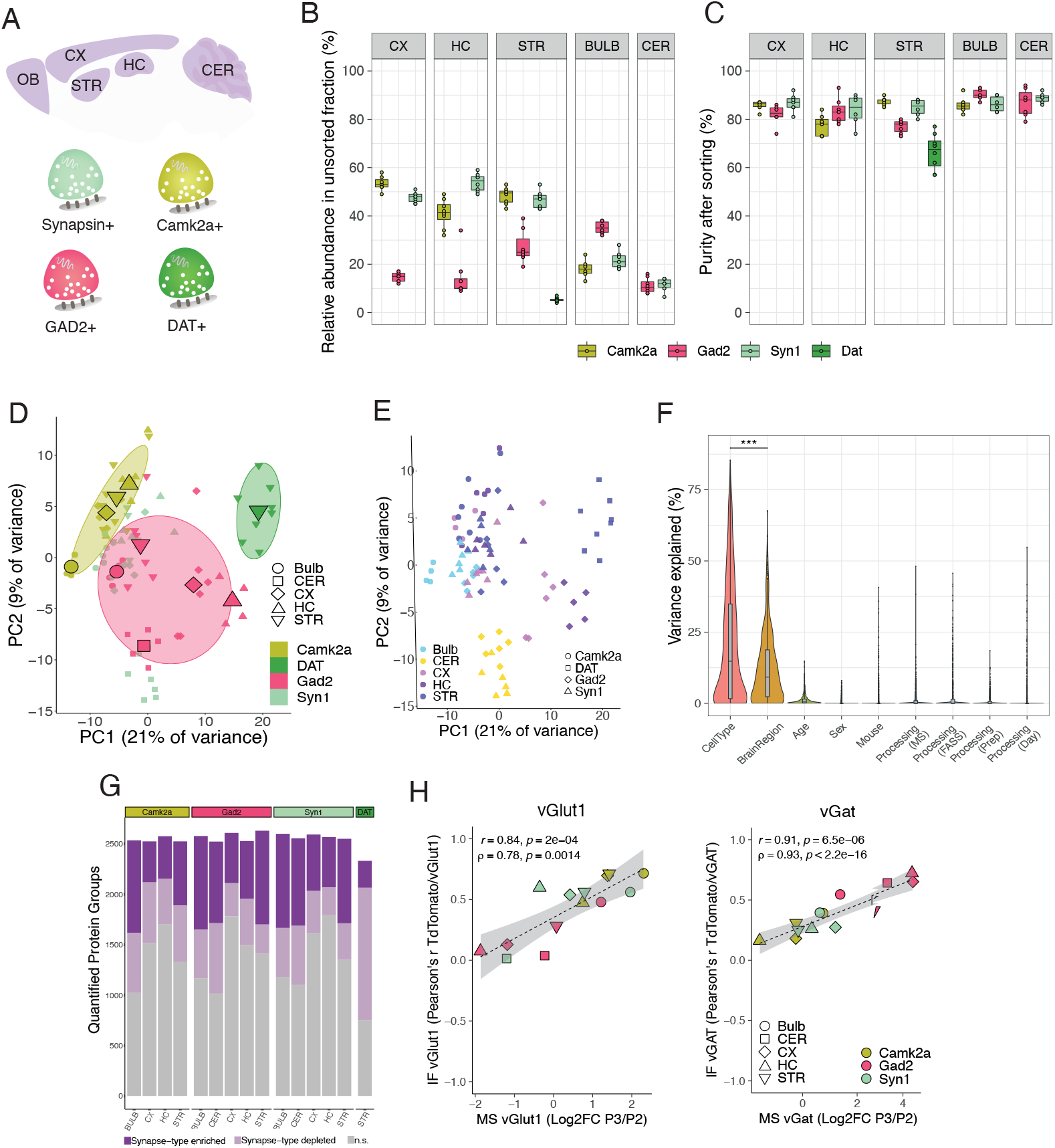
Synaptic proteomic diversity across brain areas and cell types. (A) Scheme indicating the brain areas that were microdissected from Camk2a∷SypTOM, Gad2∷SypTOM, Syn1∷SypTOM or Dat∷SypTOM mice and the resulting purified synaptosome preparations that were generated and then introduced to the pipeline. (B) Plot indicating the relative abundance of each fluorescently-labeled synaptosome type in the crude synaptosome fraction generated from the brain areas indicated (x-axis). Abundance ranged from less than 10% for Dat+ synaptosomes in the striatum crude synaptosome fraction to ~55% for the Syn1+ synaptosomes in the hippocampal crude synaptosome fraction. (C) Plot indicating the purity of each fluorescently-labeled synaptosome type from the brain areas indicated (x-axis) after FASS. The average purity of the majority of FASS synaptosomes ranged from 77% to 91%, with the exception of striatal Dat+ synaptosomes exhibiting the lowest purity of 66%. (D) Principal components analysis (PCA) in which the cell-type clusters are highlighted. Small symbols denote individual biological replicates, large symbols denote averages of each synapse subtype. Note separation of Gad2+, Camk2a+ and Dat+ cell-type clusters. (E) Principal components analysis (PCA) in which the brain regions are highlighted. Small symbols denote individual biological replicates, large symbols denote averages of each synapse subtype. Note separation of cerebellar synapse-types. (F) Violin plots depicting percentage of the variance explained by individual covariates. ***P<0.001; t test, n = 1022. (G) Number of protein groups quantified for each synapse subtype, grouped by cell types and brain regions. Shown are significantly enriched and de-enriched groups (see methods) as well as protein groups that are not significantly different between the groups. (H) Correlation between immunofluorescence and mass spectrometric measurements for vGat and vGlut1 proteins across the 15 synapse types. X-axis shows mass spectrometric measurements for vGat and vGlut1 protein, represented by the log2FC of sorted synaptosomes versus controls for each synapse type. Y-axis indicates immunofluorescence measurements for vGat and vGlut1 proteins, represented by the correlation of each synapse type’s TdTomato fluorescence intensity with vGlut1 or vGat respectively. The immunofluorescence data is described in detail in Figure S4 and 5B,C.

The fluorescently-labeled synaptosome populations that we studied originally constituted from ~5 to ~50% of a given brain region’s total crude synaptosome population **(Figure 2B).** Using the above protocol (**Figure 1A**), we purified all synapse types (but one, Dat∷SypTOM) to greater than 75% purity, on average **(Figure 2C)**. For each synapse subtype we obtained ~10M sorted (P3 gate) particles and the same number of matched control particles from at least 5 mice. We processed the sorted synaptosomes for quantitative proteome analysis using targeted feature extraction of data-independent acquisition mass spectrometry measurements (see Methods). Overall, we quantified >2’800 protein groups with a high reproducibility (median CV ~20%) for biological replicates **(Figure S8)**. In order to define the individual synaptic proteomes, we determined the complement of proteins that were quantitatively enriched in each synapse type **(see methods, Table 2, Tab1)**. In a principal component analysis the different synaptic proteomes were distinguished by both cell types **(Figure 2D)** and brain regions **(Figure 2E)**. Which feature, cell type or brain region, exerts the greatest influence on the synaptic proteomes? Overall, the cell types explained more of the total variance than the brain regions **(Figure 2F)**, and other factors like age or sex of the animals had negligible influence on the observed variance **(Figure 2F)**. In total we allocated >10’000 protein groups to the 15 synaptic proteomes **(Figure 2G)** and we identified >1’800 unique protein groups that were enriched in at least one synapse subtype **(Table 2, Tab1)**. In order to verify the obtained synaptic proteomes, we compared the enrichment of the vGat and vGlut1 proteins from mass spectrometry with those obtained by immunofluorescence on brain slices **(Figure S4)** for every synapse subtype. We found high correlation coefficients of 0.84 and 0.91 for vGlut1 and vGat respectively **(Figure 2H)**, validating the specificity of these synaptic proteomes.

To what extent do the proteins identified here represent previously known synaptic molecules? To address this, we compared the union of all the unique synapse type-enriched proteins with the synaptic protein database SynGO (Koopmans et al. 2019). Overall, there was very good agreement between the two datasets: ~80% of the identified SynGO-annotated proteins were classified as synapse-enriched in our analysis **(Figure S9)**. Furthermore, we identified >1’000 new protein groups, not previously included in SynGO, as synapse-enriched **(Table S2, Tab2)**. These novel synapse-enriched proteins are significantly associated with various disease-related pathways and underlie different cellular functions such as G-protein signaling, protein degradation or tRNA aminoacylation **(Figure S9).** When each synaptic proteome was analyzed individually, each one was significantly enriched in numerous SynGO terms, including the top-tier terms synapse, presynapse and postsynapse **(Table S2, Tab3)**. As might be expected from the localization of the SypTOM protein, we found significantly more proteins annotated as presynaptic than postsynaptic across the 15 synapse type **(Figure S9)**. While the number of new synaptic proteins identified did not differ between brain regions, we identified significantly more novel synaptic proteins at Gad2∷SypTOM synapse types as compared to Camk2a∷SypTOM **(Figure S9)**, and the most novel proteins at Dat∷SypTOM synapses, presumably reflecting a bias of the published synaptic protein literature towards excitatory synapses.

Are there protein modules that are shared amongst the different synapse types? To address this first across all types, we selected proteins that were detected in at least 14 of the 15 synapse types and performed an enrichment analysis using the SynGO database to identify functional groups that are overrepresented among the proteins that are present at most synapse types. We identified a significant enrichment of synaptic vesicle vATPases as well as proteins mediating synaptic vesicle endocytosis **(Figure S10)**. Both terms were also significantly enriched when selecting proteins that were detected in minimally 10, 11, 12 or 13 synapse type proteomes **(Figure S10)**. Next, we quantitatively compared Gad2∷SypTOM with Camk2a∷SypTOM synaptosomes and defined Gad2-enriched, Camk2a-enriched and shared protein groups in each brain region **(Figure 3A, see methods)**. This analysis reveals proteins that are quantitatively enriched at either synapse type over the other (Camk2a-enriched and Gad2-enriched) or enriched in both types over controls and not significantly different between the types (shared). In general, we observed very little overlap (~1% of all synapse-enriched proteins) between Gad2∷SypTOM enriched and Camk2a∷SypTOM enriched proteins, indicating that across all brain regions, excitatory and inhibitory synapses have defined sets of mutually exclusive synaptic proteins **(Figure 3B, see further analysis below)**. However, since Gad2-cre and Camk2a-cre label different neuron populations in different brain regions **(Figure S3-5)**, there were subsets of shared proteins in some brain regions available for analysis **(Figure 3B)**. To examine in more detail the shared proteins, we focused our analysis on the shared proteins of the Cortex (where Gad2∷SypTOM and Camk2a∷SypTOM label non-overlapping synapse populations) **(Figure S3-5)**. We found differential enrichment of proteins involved in different parts of the synaptic vesicle cycle, indicating that some protein modules of the synaptic vesicle cycle have the same proteins at both synapse types independent of neurotransmitter identity, while other protein modules show a synapse type-specific specialization in their protein composition. Within the shared proteins, we observed an overrepresentation of synaptic vesicle endocytosis and synaptic vesicle proton loading (vesicular ATPases), while exocytosis and presynaptic active zone proteins were absent from the shared fraction, but overrepresented in the Gad2-enriched and/or the Camk2a-enriched fraction **(Figure 3C)**. We constructed a protein-protein interaction network (Szklarczyk et al. 2019) using the proteins associated with the enriched terms and revealed that protein modules that represent different steps of the synaptic vesicle cycle are enriched in either shared or subtype-specific synaptic proteins **(Figure 3D)**. These analyses highlight endocytosis and vesicular ATPases as generic synaptic protein modules that are used by synapses independent of neurotransmitter type, while the exocytosis machinery, the presynaptic active zone, trans- and postsynaptic protein modules are composed of different sets of proteins for different synapse types.

**Figure 3.**
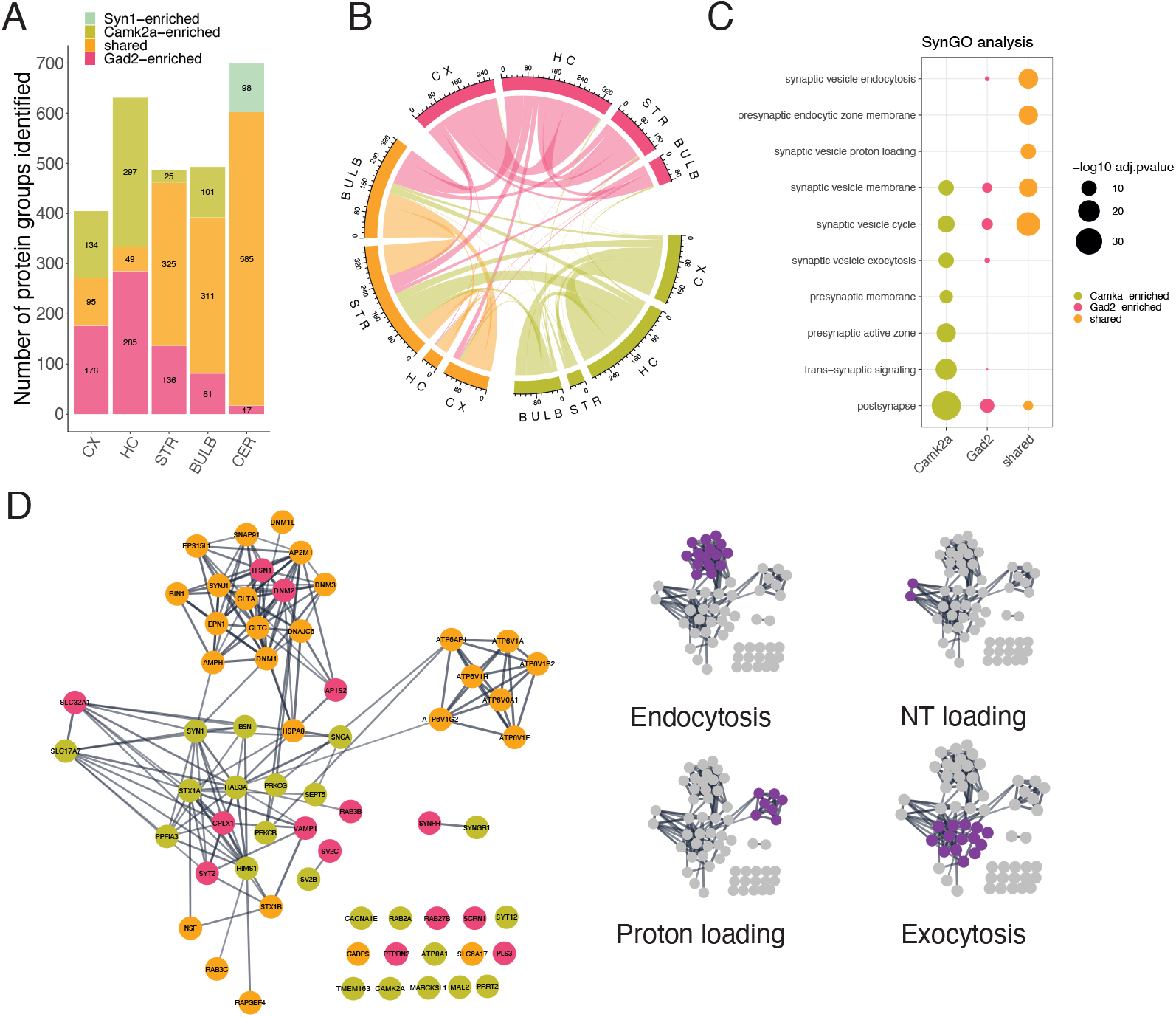
Synaptic proteome commonalities and differences. (A) Barplot showing proteins significantly enriched in the direct quantitative comparison of Camk2a+ vs. Gad2+ synaptic proteomes for each brain region. Shared proteins are defined as significantly enriched in both Camk2a+ and Gad2+ versus control synaptosomes and not significantly different between Gad2+ and Camk2a+. Note increased number of shared proteins in brain regions where Camk2a-cre and Gad2-cre label exclusive as well as overlapping populations and smaller numbers of shared proteins in regions where Camk2a-cre and Gad2-cre label mutually exclusive cell types (Figure S3-5). The cerebellum lacked detectable TdTomato signal in the Camk2a∷SypTOM mouse so the Syn1+ proteome was used for the Gad2+ comparison here. (B) Chord diagram of intersections between the groups defined in A; with the 3 colors representing Camk2a+-enriched (green) Gad2+ enriched (red) or enriched in both (yellow). The arcs indicate overlapping proteins between the two connected groups. This analysis allows one to better observe the extent of overlapping proteins between distinct brain regions. Note that there are few intersections (~1% of synapse-enriched proteins) between Camk2a and Gad2 relative to Gad2 with shared and Camk2a with shared meaning that across all brain regions, excitatory and inhibitory synapses have defined sets of mutually exclusive synaptic proteins. (C) Dotplot of SynGO analysis results for shared and cell-type specific enriched proteins. Depicted are selected significantly enriched SynGO terms of Camk2a-enriched, Gad2-enriched and shared groups from cortex. (D) Protein interaction network of synaptic vesicle cycle proteins for cortical Camk2a, Gad2 or shared-enriched groups. Proteins with SynGO annotation for the synaptic vesicle cycle are displayed. Edges represent a stringdb score >0.7 (high confidence) (Szklarczyk et al. 2019). Proteins that are associated with the significantly enriched terms “synaptic vesicle endocytosis”, “synaptic vesicle exocytosis” and “synaptic vesicle proton loading” (data shown in C) are indicated on the left.

Are there specific protein modules that are associated with synapses that use different neurotransmitters or are associated with different cell types or brain regions? To address this we used a protein-protein weighted network correlation analysis (WGCNA) (Langfelder and Horvath 2008) to identify protein modules that show correlated abundance patterns across the 15 synapse types. We first asked a simpler question: do proteins of the same complex or functional unit exhibit correlated expression levels? We found a significantly higher median correlation for proteins that are subunits of the same protein complex (pearson’s r = 0.65) (Giurgiu et al. 2019) as compared to random protein pairs (pearson’s r = 0.03) **(Figure 4A)**. For example, the proteasome subunits Psma1 and Psma7 exhibited highly correlated abundance (r = 0.82); similarly, vGat and Gad2 (the vesicular GABA transporter and the essential synaptic GABA synthesis enzyme) were correlated with a near perfect coefficient of 0.91. We next constructed a protein-protein correlation network using all synapse-enriched proteins and identified 14 protein modules using WGCNA **(Figure 4B, Table S3)**. We discovered that the resulting protein network featured two main opposing clusters, defining two highly connected protein communities. To identify the nature of these protein communities, we correlated all protein modules with traits of the synapse subtypes, including immunofluorescence for vGat and vGlut1 **(Figure 2H and S6)**, cell types and brain regions. While most brain regions and the cell type Syn1 showed no correlation with any protein module, we identified three inhibitory protein modules that were significantly correlated with vGat and three excitatory protein modules that were significantly correlated with vGlut1 **(Figure 4C)**. We found that both the excitatory and inhibitory modules were located at the center of each protein community, and that each community was characterized by high correlation or anticorrelation with vGat or vGlut1 **(Figure 4D, Table S3)**. In summary, we present a protein-protein correlation network that features >1’500 synapse-enriched proteins and reveals two distinct protein communities that represent the proteomes associated with excitatory or inhibitory neurotransmitters across distinct cell types and brain regions.

**Figure 4.**
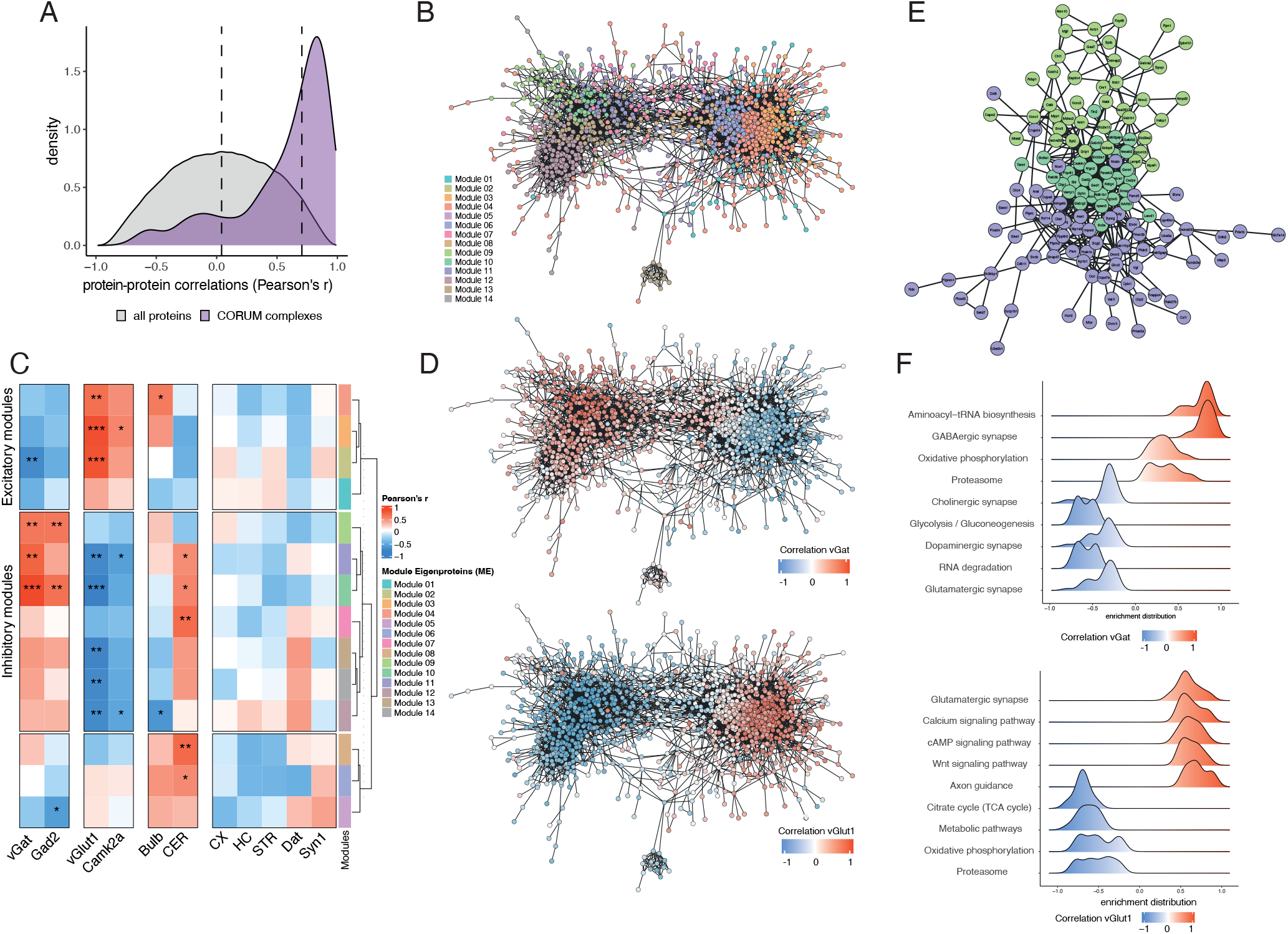
The synaptic protein-protein correlation network reveals protein communities. (A) Density plot of pairwise protein-protein abundance profile correlations (pearson’s r) for protein pairs that are annotated members of the same protein complex (purple) (Giurgiu et al. 2019) and for random protein pairs (grey) as control. Proteins of the same complex exhibited a highly co-regulated abundance profile across the 15 synapse types, while random protein pairs showed no correlation on average. Dashed lines denote median values for each group. (B) Community network of protein-protein correlations. The network represents a visualization of the adjacency matrix used for WGCNA. The nodes are synapse-enriched proteins and they are connected by edges that represent the abundance profile correlation of the two nodes they connect. Specifically, edges represent adjacency based on biweight midcorrelation and are filtered for weights >0.3, meaning negative and low correlations are not considered for visualization of the network. Protein nodes are colored according to their associated protein module. (C) Heatmap of module correlations with synaptic traits. Protein module eigenproteins are correlated with the following traits of the 15 synapse types; cell-type, brain region and immunofluorescence for vGat and vGlut1. Protein module eigenproteins are protein abundance profiles that are representative for the proteins in their module (specifically, the first principle component of the module). Significant correlation of a protein module with a synaptic trait suggests that the proteins in that module are important for the trait. *P<0.05; **P<0.01; ***P<0.001; pearson correlation.

Can we identify the key proteins for the synaptic proteomes of glutamatergic and GABAergic synaptic proteomes based on the network topology? We identified 188 and 315 proteins that are significantly correlated with vGat or vGlut1, respectively, with correlation coefficients ranging from moderate (~0.5) to very high (~0.95) **(Table S3)**. Proteins with high correlation coefficients were also among the most connected nodes within each community **(Figure 4D)**. We inspected the network topology of the protein modules with a significant correlation for vGat and found that the core module (Module 10) contained established inhibitory marker proteins, including Gad-1, Gad-2, Gaba transporter-1, Neuroligin-2, multiple GABA-A receptor subunits (alpha-1, gamma-2, beta-2 and beta-3) and the inhibitory postsynaptic scaffolding protein gephyrin **(Figure 4E)**. Notably, Module 10 contained proteins from both the pre- and postsynaptic compartments, indicating a tight co-regulation of synaptic architecture across the synapse. The Module 09, in contrast, was characterized by moderate but significant correlation with vGat. In this module we detected proteins that have been previously associated with subtypes of GABAergic neurons or synapses, for example Syt-2 (Sommeijer and Levelt 2012), Lamp-5, Cannabinoid-receptor-1 or GABA-A receptor subunit alpha-2 (Zeisel et al. 2018) **(Figure 4E)**. In total, we identified 130 novel synaptic proteins that correlate significantly with vGat and were not previously recognized as synaptic by SynGO. In the vGat core protein module we identified, for example, IgLON5, a cell adhesion protein implicated in a specific anti-IgLON5 neurodegenerative autoimmune disease (Madetko et al. 2022).

In order to validate the network topology of the GABAergic proteome, we constructed a protein-protein interaction network of the three inhibitory protein modules using a protein interaction database and found that the proteins from the core module (Module 10) accounted for the majority of hubs at the center of the network **(Figure S11)**. In conclusion, we present a protein-protein correlation network that represents the diversity of the GABAergic synaptic proteome, revealing many novel synaptic proteins and highlights core proteins that are strongly associated with vGat across synapses from different brain regions and cell types, as well as moderately associated proteins that may modulate specific synapse subtypes.

The excitatory synapse protein community was well-correlated with vGlut1 fluorescence **(Figure 4D)**; correspondingly, we found the vGlut1 protein was among the most connected nodes and there was no significant correlation with the vGlut2 protein. As such, the identified protein community was therefore specific for the most abundant (vGlut1+) excitatory synapse type. Overall, we detected many more proteins in the vGlut1+ network, when compared to the vGat+ network. Presumably, this is due to the elaborate postsynaptic density complex and spine architecture present at excitatory synapses that is typically absent at inhibitory synapses. Consistent with this idea, we detected many proteins from established excitatory postsynaptic protein families within the vGlut1 protein community, including the Shank family (Shank1/2/3), the Camk family (Camk4/1d/2d/2b/v/2a), the Dlg family (Dlg1/2/3/4, Mpp2/3 and Dlgap1/2/3/4), the glutamate receptor subunits (Gria1/2/3/4, Grin1/2b, Grik2), the Lrr family (Lrrc7/57/4) and protein phosphatases (Pp2r5a/3ca/1cb/5c/1ca/3cb/3ra) **(Figure S11)**. We also identify numerous proteins not previously associated with glutamatergic synapses. For example, we found Fbxl16, an F-box protein of E3-ubiquitin ligase with unclear synaptic function (Honarpour et al. 2014; Kim et al. 2021), exhibited the strongest correlation with vGlut1 across brain regions. In total, we identified 158 novel synaptic proteins that correlate significantly with vGlut1 and were not previously recognized as synaptic by SynGO.

We next asked whether there are pathways or biological functions enriched specifically at vGat versus vGlut1 protein communities. To address this, we performed gene set enrichment analysis using the ranked protein correlations with vGat and vGlut1 immunofluorescence, respectively. As expected, we found significant enrichment for proteins associated with postsynaptic signaling pathways and dendritic spines within the vGlut1 community **(Figure 4F and S11)**. In the vGat protein community, we found significant enrichment for proteins from the GABA receptor complex, but also proteins with functions not previously recognized as enriched at inhibitory (versus excitatory) synapses: aminoacyl-tRNA synthetases, proteasome subunits and mitochondrial proteins **(Figure 4F and S11)**. Taken together, the above analyses define a roadmap for how molecular systems-biology analysis of synaptic proteomes can be used to identify the key protein modules that underlie synaptic traits. The resulting protein-protein correlation network comprising >1’500 synaptic proteins revealed shared and specialized synaptic protein modules according to neurotransmitter identity.

In contrast to glutamate and GABA, dopamine represents the class of modulatory neurotransmitters. Dopaminergic synapses have been intensively studied in the context of midbrain dopaminergic neurons that project to the striatum, which are critically important for reward processing and movement control (Liu, Goel, and Kaeser 2021), and their degeneration is a main hallmark of Parkinson’s disease pathophysiology (Poewe et al. 2017). We conducted an in-depth analysis of the modulatory synaptic proteome of dopaminergic terminals in the striatum. To verify the expression of the presynaptic fluorophore in the Dat∷SypTOM mice we immunostained brain sections (**Figure 5A**) and examined the coincidence of the TdTomato signal with different markers. As expected, there was a prominent TdTomato signal in the striatum and we found high correlation with Tyrosine Hydroxylase immunoreactivity (**Figure 5B and C**). Comparing striatal Dat∷SypTOM synaptosomes with unsorted control synaptosomes, we identified 267 significantly enriched proteins in the striatal dopaminergic proteome **(Table S2, Tab 1)**. Are there proteins that are specific for the dopaminergic proteome, and which proteins might be shared between dopaminergic and other synapse types? To address this we identified proteins that were either enriched or de-enriched in striatal dopaminergic synapses as compared to the other 14 synapse types. Among the differentially enriched proteins we found dopaminergic marker proteins, for example, Maoa and Aldh1ha1, but also proteins that have not been previously associated with dopaminergic terminals (**Figure 5D and S12)**. For example, we identified Oxr1, Oxidation resistance protein 1, as ubiquitously present in Camk2a, Gad2 and Syn1 synapse types but depleted from dopaminergic terminals (**Figure 5D**). Oxr1 controls sensitivity to oxidative stress (Oliver et al. 2011; Williamson et al. 2019) and as such, the absence of Oxr1 in dopaminergic synapses might confer susceptibility to oxidative damage, a major contributor to dopamine neuron degeneration observed in Parkinson’s disease. Another example is the mitogen-activated protein kinase Erk1 (Mapk3), which was specifically enriched at dopaminergic synapses (**Figure S12**) and has been linked to Parkinson’s disease via multiple cellular processes (Bohush, Niewiadomska, and Filipek 2018). Next, we quantitatively compared the striatal Dat∷SypTOM proteome to the striatal Syn1∷SypTOM proteome (**Figure 5E**) and performed gene set enrichment analysis comparing mutually enriched with Dat∷SypTOM specific proteins (**Figure 5F**). Relative to striatal Syn1∷SypTOM synapses, the Dat∷SypTOM synaptic proteome was highly enriched in key proteins involved in dopamine biosynthesis, trafficking and degradation (**Figure 5E and G**). The proteins that were significantly enriched in both groups included vesicular ATPases, proteasome subunits and endocytic proteins (**Figure 5E and Figure S12**). While we identified four vATPases within the shared group (Atp6v1a/h/f/e1), Atp6v1g1 was significantly enriched at dopaminergic synapses, indicating an association with dopaminergic synaptic vesicles (**Figure 5G).** We identified six proteasome subunits as shared, and two proteasome subunits that constitute the modulatory PA28 complex (Psme1/2) as enriched at dopaminergic synapses. The PA28 complex associates with the immunoproteasome, which generates MHC peptides in myeloid cells, and has been shown to facilitate the degradation of oxidized proteins, thereby contributing to adaptation and tolerance of oxidative stress (Pickering et al. 2010). We validated the presence of the Psme1 protein and proteasome activity in striatal synaptosomes **(Figure S12)** but found no evidence for presence of immunoproteasome subunits. Instead, we found Psme1 in complex with the standard proteasome in cultured neurons (**Figure S12)**. Together, these findings suggest a role for PA28 in a complex with the constitutive proteasome at dopaminergic synapses. The above data identify the synaptic proteome of striatal dopaminergic terminals and highlight similarities and specializations in synaptic proteome architecture.

**Figure 5.**
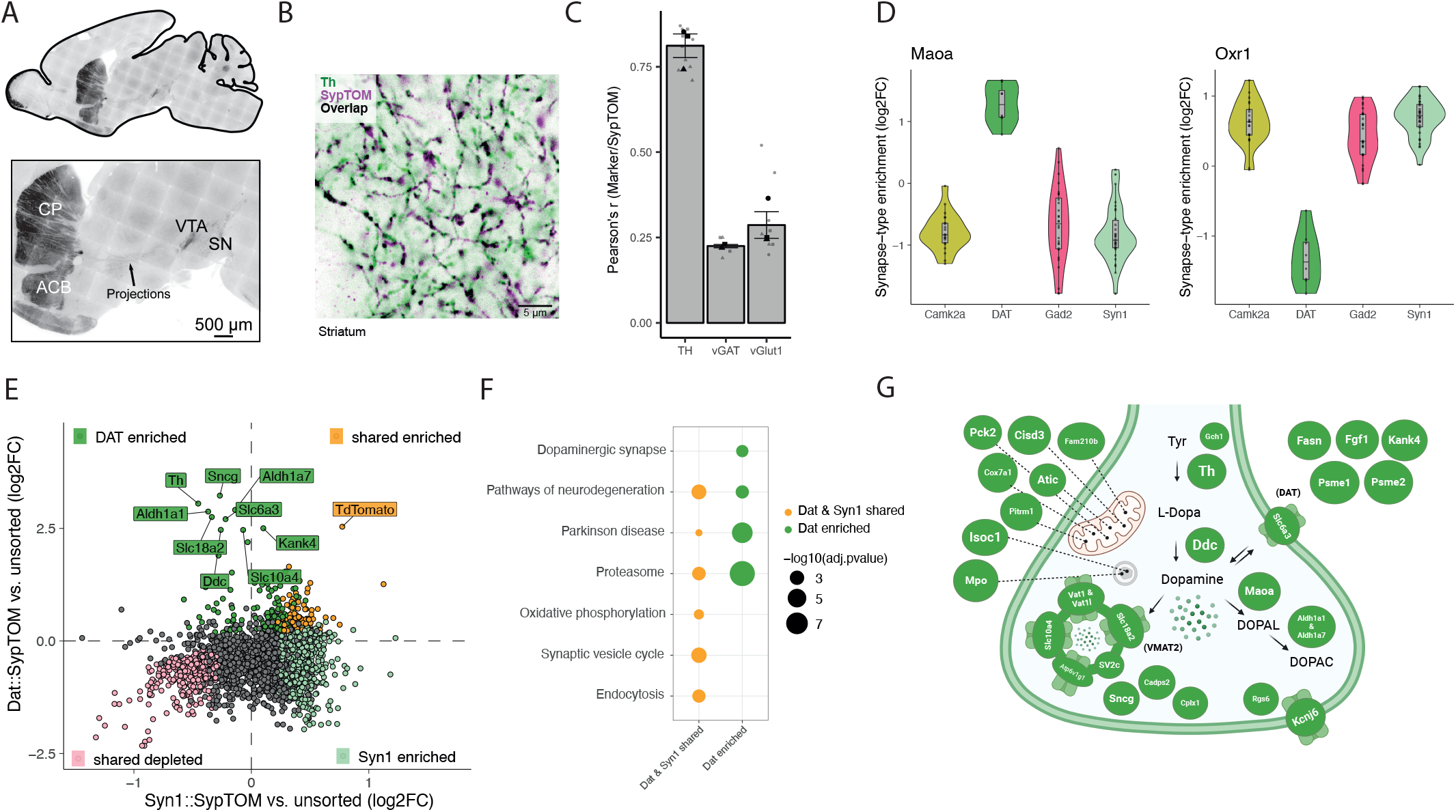
The dopaminergic synaptic proteome. (A) Representative low (upper) and high (lower) magnification images of an immunostained brain section from a Dat∷SypTOM mouse showing TdTomato present in the nigrostriatal pathway. Fluorescent signal is detected in the cell bodies of the ventral tegmental area (VTA), substantia nigra (SN) and their associated projections to striatal areas caudate putamen (CP) and nucleus accumbens (ACB). (B) Representative image depicting overlap between tyrosine hydroxylase (Th) immunoreactivity with TdTomato fluorescence at high magnification in the striatum of a Dat∷SypTOM mouse. (C) Analysis of data shown in B supplemented by correlation of TdTomato immunoreactivity with vGlut1 and vGat in the striatum of Dat∷SypTOM mice. n = 2-4 animals, 2-4 images per mouse, larger data points represent biological replicates, while smaller points depict individual images. Error bars signify the standard error of the mean. (D) Violin plots for two representative proteins showing specific enrichment (Amino oxidase A, Maoa, a marker for dopaminergic neurons) or specific depletion (Oxidation resistance protein 1, Oxr1) in dopaminergic synaptic terminals compared to all other synapse types. (E) Scatter plot comparing the differential enrichment of proteins in the Dat+ and Syn1+-synaptosomes to their striatal control synaptosome precursor populations. Colors indicate proteins that are significantly enriched in Dat+-sorted synaptosomes (green), Syn1+-sorted synaptosomes (pale cyan), significant in both populations (orange) and significantly de-enriched in both (pale pink). Note the specific enrichment of labeled dopaminergic marker proteins. (F) Dotplot of selected significantly enriched pathways (KEGG) of GSEA comparing exclusively dopaminergic synapse-enriched proteins with shared enriched proteins between dopaminergic and all striatal synapses (Syn1+) (F) Scheme showing selected top-enriched proteins (Dat+ compared to unsorted controls or Syn1+) within the presynaptic terminal.

We next asked whether different synapse types characterized by the same neurotransmitter class exhibit diversity in their synaptic proteomes. The neurons that use GABA as a neurotransmitter exhibit strong morphological diversity and differ in the location of their cell bodies and their synaptic contacts on other cells (Huang and Paul 2019), as well as their transcriptomic profiles, connectivity patterns and firing properties (Fishell and Kepecs 2020). Using cre-driver lines for the main subclasses Parvalbumin-(PV), Somatostatin-(SST) and vasoactive intestinal peptide-(VIP) neurons we targeted synapses arising from the three largest groups of molecularly-defined cortical GABAergic neuron subclasses (Taniguchi et al. 2011) (**Figure 6A**) and compared them to the cortical Gad2∷SypTOM synaptic proteome using the Gad2-cre driver line. We verified the co-localization of TdTomato signal with immunoreactivity for established markers for each of the three subtypes in brain slices (**Figure S13**). Cortical interneurons, however, are very sparse, making their synaptic proteomes very challenging to purify and analyze. For example, all GABAergic neurons make up just ~20% of neurons in the cortex and Gad2+ synaptosomes represent about 15% of the total synaptosome population (**Figure 6B**), suggesting that sub-types of GABAergic synapses would be challenging to purify. Indeed, we found that VIP+ synaptosomes represented just ~0.5% of all particles in a cortical synaptosome fraction, while PV+ and SST+ synaptosomes represented approximately 3% and 5% (**Figure 6B**). Therefore, we downscaled and optimized our sample processing to accommodate an input as low as 2 Mio synaptosomes while retaining synaptic proteome depth (**see methods**). After sorting, we were able to achieve greater than 80% purity for all 3 cortical inhibitory synapse types (**Figure 6C**). Using this optimized workflow, we quantified >2500 protein groups in total and identified almost 600 unique proteins enriched in at least one cortical interneuron subtype, thereby outlining the overall cortical inhibitory synaptic proteome. We found distinct synaptic proteomes for the three subtypes with 325, 234 and 369 significantly enriched proteins detected respectively for PV+, SST+ and VIP+ synapses (**Figure 6D, Table S4, Tab 1**). All proteomes were significantly enriched with GABAergic synaptic proteins (**Figure S14).** The three inhibitory synaptic proteomes were clearly separated in a principal component analysis (**Figure 6E**) and showed a type-specific signature in their proteome composition (**Figure 6F**). The union Gad2 proteome was closest to the most abundant type SST+ in PCA space. We searched for canonical markers of each inhibitory synapse type and found, as predicted, the expected enrichment for VIP, PV, and Calbindin proteins in the VIP+, PV+ and SST+ synaptic proteomes (**Figure 6G**). The remaining markers, Lamp5 and Scng, distinguished a 4^th^ and 5^th^ type of inhibitory neuron (Huang and Paul 2019; Taniguchi et al. 2011) and, appropriately, were most enriched in the Gad2+ cortical synaptosomes (**Figure 6G**).

**Figure 6.**
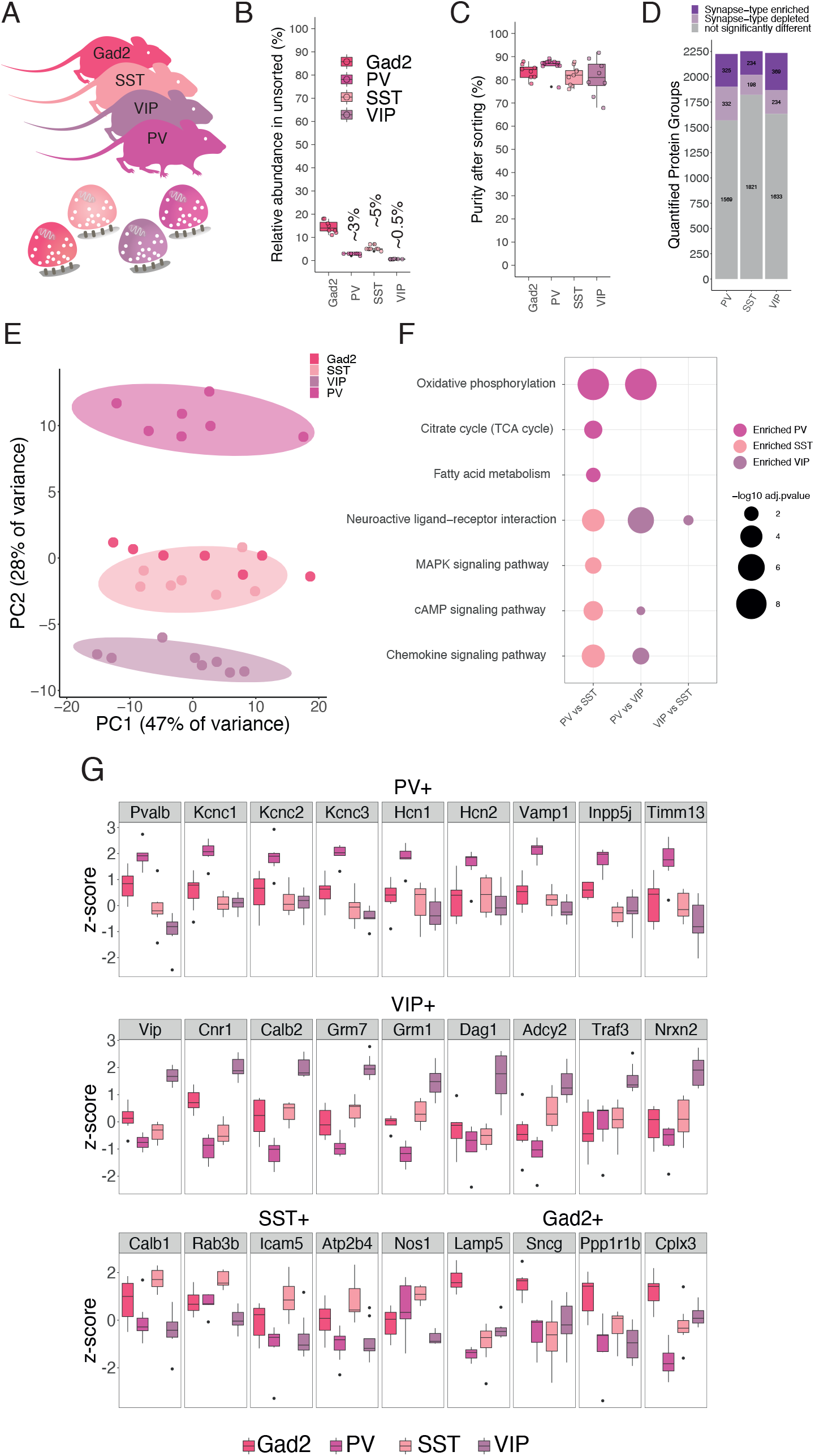
Proteomic diversity of Gad2, Parvalbumin, Somatostatin and Vasointestinal active peptide synapses. (A) Scheme showing the different mouse-lines from which cortical synaptosomes were prepared and the resulting purified synaptosome populations. (B) Plot indicating the relative abundance of each fluorescently-labeled synapse type in the crude cortical synaptosome fraction. As predicted, abundance was very low for all interneuronal populations. (C) Plot indicating the purity of each fluorescently-labeled cortical interneuron synaptosome type after FASS. For all types, the average purity exceeded 80%. Purity is assessed by re-analysis of the sorted fraction by synaptosome flow cytometry. (D) Number of protein groups quantified for each cortical interneuron synapse subtype. Shown are significantly enriched and de-enriched groups (see methods) as well as protein groups that are not significantly different between the groups. (E) PCA of synaptic proteomes from cortical inhibitory subtypes. (F) Dotplot of selected significantly enriched pathways (KEGG) of GSEA comparing cortical interneuron types directly against each other. Analysis is based on protein lists ranked by log2FC of the indicated synapse types in the X-axis. (G) Boxplots for representative proteins that show specific enrichment in the indicated cortical interneuron subtype.

Do the above identified cortical interneuron proteomes reflect the vGat protein communities defined by the protein-protein correlation network **(Fig 3)**? We identified 89% of the core vGat module proteins enriched in at least one cortical interneuron subtype and 63% were enriched in all three types, indicating a high degree of agreement between the two approaches. Of all proteins in the vGat community (defined by significant correlation with vGat), we identified 73% that were also enriched in the overall cortical inhibitory proteome, including 83 proteins not previously annotated as synaptic in SynGO. We next asked whether the developmental origin of the cell type is reflected in the synaptic proteome. While PV and SST neurons arise from the medial ganglionic eminence (MGE), VIP neurons originate from the caudal ganglionic eminence (CGE) (Kepecs and Fishell 2014). Previous studies showed that transcriptomes of cortical inhibitory neurons (BRAIN Initiative Cell Census Network (BICCN) 2021; Zeisel et al. 2018; Fishell and Rudy 2011) cluster according to their progenitor domain. In contrast, principal component analysis revealed that the SST+ and VIP+ synaptic proteomes cluster closer together than SST+ and PV+ proteomes (**Figure 6E**). Consistent with this, we identified only 33 proteins that were significantly different between the SST+ and VIP+, but 128 and 318 between PV+ and SST+ or VIP+ respectively (**Table S4, Tab 2,3 and 4**). Together, these analyses indicate that cortical inhibitory synaptic proteomes are predominantly shaped by other factors than the developmental origin of their presynaptic cell type.

Can we identify proteins that relate to the functional differences observed in these three inhibitory synapse types? PV+ neurons are characterized by fast spiking and corresponding high energy demands. We found that the voltage-gated potassium channels Kcnc1, Kcnc2 and Kcnc3 (Kv3.1/2/3) were among the most enriched PV+ synaptic proteins proteins (**Figure 6G**), potentially explaining the ability of PV cells to fire at high rates (Kaczmarek and Zhang 2017). Subcellular patch-clamp recordings have demonstrated that axonal hyperpolarization-activated cyclic nucleotide-gated ion (HCN) channels counter-balance the activity-dependent hyperpolarization during high-frequency firing in PV neurons (Roth and Hu 2020). Correspondingly, Hcn1 and Hcn2 were highly enriched in the PV+ synaptic proteome (**Figure 6G**). Consistent with their high firing rate and associated energy demands, the PV+ proteome was also significantly enriched in mitochondrial proteins involved in oxidative phosphorylation (**Figure 6F**). Additionally, we detected a number of synaptic vesicle-associated proteins (such as Syt2, Vamp1 or Cplx1) and the cell adhesion proteins Cntnap4 (Karayannis et al. 2014) and Hapln4 as specifically enriched at PV+ synaptic proteomes (**Figure 6G, Table S4**).

Cortical VIP neurons mainly inhibit SST and PV interneurons and thereby constitute a critical component of cortical disinhibitory circuits. They are characterized by heterogeneous firing patterns, suggesting diverse functional roles. In comparison to the PV+ proteome, the VIP+ synaptic proteome was enriched for neuroactive ligand-receptor interactions and downstream signaling molecules (**Figure 6F,G**). One of the most enriched proteins was Cannabinoid receptor 1, Cnr1, and many proteins involved in downstream G-Protein signaling were enriched as well, including Rgs6, Kcd12, Adcy2, Gng2 and Gnai2 (**Figure 6G, Table S4**). We also detected a strong enrichment of glutamate G-Protein coupled receptors, the metabotropic glutamate receptors (excitatory Grm1 and inhibitory Grm7), and to a lesser extent the ionotropic glutamate receptor subunits Grik2 and Gria1/4 but not Gria2/3 (**Figure 6G, S14 and Table S4**). Furthermore, we found a number of cell-adhesion proteins specifically enriched at VIP+ synapses (Nrxn1,Nrxn2, Nlgn3, Dag1 and Igsf8), calcium-binding proteins (Calb2, Necab2), calcium channels (Cacna1b, Cacna2d3) and synaptic-vesicle associated proteins (Sh3gl3, Rph3a, Rab3c, Synpr and Syn3) (**Figure 6G, S13 and Table S4**). In contrast to PV+ and VIP+, there were few proteins (Calb1, Rab3b, Icam5, Nipsnap3b, Atp2b4 and Nos1) that distinguished SST+ synaptic proteomes from both of the other two types (**Figure 6G**). This reflects the current view that SST neurons represent a diverse group with substantial differences in morphology and physiology (Urban-Ciecko and Barth 2016). One SST+ enriched synaptic protein was Atp2b4, which physically interacts with nitric oxide synthase (Gillespie et al. 2022), Nos1, also highly enriched at SST+ synaptic proteomes compared to Gad2+ **(Figure 6G).** These findings align with previous reports classifying Nos1-expressing neurons as a SST subtype (Kubota et al. 2011). Finally, we identified few proteins (Ppp1r1b and Cplx3) that were enriched in the overall Gad2 synaptic proteome over the three inhibitory subtypes, presumably because they are specific for one of the other main cortical interneuron subtypes characterized by Lamp5 or Scng expression **(Figure 6G)**. Overall, we identified ~600 unique proteins that define the cortical interneuron synaptic proteome, allocated them to the three main subclasses and highlighted specialized groups of proteins that relate to their established functional properties.

## Discussion

Synapses come in all shapes and sizes, and their plasticity in structure and function is crucial for correct wiring of neuronal circuits, which in turn encode the cognitive capabilities of an organism. While there is evidence for substantial structural and functional diversity of synapses (O’Rourke et al. 2012), the underlying diversity in synaptic molecular architecture is much less understood. A detailed understanding of synapse proteome diversity allows us to link the molecular architecture of the synapse to structure and function. Here, we developed and optimized FASS (Biesemann et al. 2014) in combination with mass spectrometry for system-wide analysis of the proteomic landscape of synaptic diversity across 18 distinct synapse types defined by cell-type and brain region. We use a conditional SypTOM mouse line crossed with different cre-driver lines to achieve cell-type specific labeling of synaptosomes. We optimized the interface between FASS and mass spectrometry to enable deep coverage of very small numbers of synaptosomes, enabling us to profile the proteomes of rare synapse types with deep coverage at scale. Finally, we use a weighted protein co-expression network analysis (Langfelder and Horvath 2008) to identify the key protein modules in the network that are correlated with external synaptic traits. Altogether, we defined a roadmap for a molecular systems-biology analysis of synaptic proteomes, which we used to identify >1’800 unique synapse-enriched proteins. These proteins provide the building blocks for 18 synapse-type specific proteomes and we revealed that synaptic proteins form protein communities characterized by varying degree of association to vGat and vGlut1.

Our resource of synaptic proteomes departs from previous studies of synaptic proteomes in many respects. First, we used FASS (Biesemann et al. 2014) because it enables analysis of synaptic proteomes originating from *in vivo* brain structures and covers all synaptic compartments, including pre-, post and trans-synaptic proteins. Second, we cover scarce synapse types not previously amenable to purification, with VIP+ synaptic proteomes being the rarest type included in this study, representing <1% of the total cortical synapse pool. Third, the increased throughput enabled us to include many biological replicates per synapse type (>5), this increased the depth of the synaptic proteomes we obtained and allowed us to define synapse-enriched proteins in a purely data-driven fashion using linear-mixed effects models (Choi et al. 2014), without the requirement for external lists or prior knowledge. Fourth, the increased throughput allowed us to compare proteome diversity across 15 synapse types, while all previous studies were limited to one or two synapse types (Uezu et al. 2016; Spence et al. 2019; Loh et al. 2016; Apóstolo et al. 2020; Biesemann et al. 2014; Paget-Blanc et al. 2022; Hobson, Choi, et al. 2022; Wilhelm et al. 2014; Boyken et al. 2013; Takamori et al. 2006; Roy et al. 2018). The depth and breadth of our proteome coverage also allowed us to use more sophisticated systems-biology data analysis approaches as compared to binary differential expression analysis. The resulting protein-protein correlation network thus has increased confidence because it is based on a very large dataset. Furthermore, the resulting protein-protein correlation network associates previously understudied proteins to protein modules and/or synaptic traits. Importantly, this resource can not only be mined for biological insights, but also to generate hypotheses for physical or functional protein-protein interactions, or for the identification of synapse types that break the correlation between co-regulated proteins, which could indicate synapse-type specific protein complex composition (Lapek et al. 2017). Finally, our resource enables a direct comparison of different synapse types, in contrast to comparisons between synapses and other subcellular compartments like the soma (Hobson, Choi, et al. 2022). Therefore, proteins that we identify enriched at synapse type A over type B, or correlated with vGat or vGlut1, might be present in neuronal dendrites, axons or somata to varying degrees. The observed enrichments or de-enrichments are thus presumably a function of protein abundance and specific subcellular targeting or exclusion of proteins from synapses. For example, proteins that showed cell-type specific expression but broad localization throughout the cell, like the parvalbumin protein, were identified as specifically enriched when compared to other synapse types. In contrast, specific exclusion or synaptic recruitment could lead to synaptic enrichment or de-enrichment despite comparable average protein abundance in different neuron types. For example, Oxr1 is specifically enriched at many synapse types but depleted from dopaminergic terminals, the corresponding mRNA however was found at comparable levels throughout many neuron types, including midbrain dopaminergic neurons and GABAergic neurons in the striatum (Zeisel et al. 2018).

We identified hundreds of unique proteins with previously undetected known synaptic localization and we allocated thousands of proteins to different subtypes. We find vATPases and synaptic vesicle endocytosis proteins as commonly shared synaptic modules, and the presynaptic active zone, exocytosis machinery, trans-synaptic and postsynaptic elements as hotspots for synaptic proteome specialization. Using a guilt-by-association approach we found that synaptic proteins form communities that correlate with vGat and vGlut1. Intriguingly, we discovered many proteins at the core of the vGat and vGlut1 protein communities which were not previously recognized as synaptic. The vGat protein community further revealed an intriguing enrichment of unanticipated functional protein groups: we identified proteasome subunits, mitochondrial proteins as well as tRNA synthetases preferentially enriched over the vGlut1 synaptic community. The presence of mitochondrial proteins in the vGat community presumably relates to increased energy demands, analogous to the finding of increased mitochondrial proteins at the PV+ synaptic proteome. While moonlighting functions have occasionally been ascribed to tRNA synthetases (M. Guo and Schimmel 2013), it is possible that their enrichment relates to the co-enrichment of the proteasome, and they scavenge amino acids that originate from local proteasomal degradation. In contrast to vGat and vGlut1 synapses, dopaminergic neurotransmission is modulatory, and we provide in-depth analysis of striatal dopaminergic synapses. Our analysis reveals 267 proteins significantly enriched at dopaminergic terminals, which is an almost 5-fold greater depth compared to a recent proteome analysis of dopaminergic terminals (Paget-Blanc et al. 2022). We compared the dopaminergic proteome to the other 14 synaptic proteomes and specifically against the other synaptic proteomes of the striatum. Besides proteins involved in dopamine biosynthesis, degradation and transport, we identified the absence of Oxr1, a protein that protects from oxidative damage and might render dopaminergic neurons particularly susceptible to oxidative stress (Jiang et al. 2019; Williamson et al. 2019). Furthermore, we find enrichment of an alternative proteasome cap (PA28) that stimulates proteasomal degradation of peptides. While the PA28 cap was predominantly studied in association with the immunoproteasome, our findings suggest that PA28 cap associates with the constitutive proteasome in neurons. Finally, we identified ~600 unique proteins that define the synaptic proteomes of the main cortical interneuron subclasses. In contrast to the transcriptomes of cortical interneurons, we find that the synaptic proteomes do not cluster according to their progenitor domain, indicating that synaptic proteomes are shaped by other factors than the developmental origin of their presynaptic cell type. We reveal type-specific signatures in the cortical interneuron proteomes that relate to their established functional properties. For PV+ synapses, which are characterized by high firing frequencies, we identify specific enrichment of mitochondrial proteins, voltage-gated potassium channels and hyperpolarization-activated cyclic nucleotide-gated ion (HCN) channels. VIP+ synapses show enrichment of G-protein signaling, prominently Cannabinoid receptor 1, metabotropic glutamate receptors and associated downstream effector proteins. We detected the differential distribution of individual cell adhesion molecules (e.g. Nrxn1, Nrxn2, Cntnap4, Hapln4 and Icam5) suggesting a role in the distinct connectivity patterns observed for the interneuron subtypes. Intriguingly, we also observed differential abundance of proteins in the same family. Besides Calb1 (SST+) and Calb2 (VIP+), we identified Cplx1 (PV+) and Cplx3 (Gad2+) as well as Rab3b (SST+) and Rab3c (VIP+) specifically enriched at different interneuron synapse types. The synaptic proteomes of PV, SST and VIP neurons reported here will serve as a rich resource for the neuroscience community to map molecular architecture to synapse physiology, morphology and connectivity.

This study also opens the door for further molecular systems-biology analyses of synapses. The number of synaptic molecules that contribute to multiple molecular pathways running in parallel generates a complexity that has been beyond comprehension (Südhof 2018). Cellular systems can perform elaborate computations and integrate different stimuli with non-trivial relationships (Nandagopal et al. 2018; Antebi et al. 2017). Holistic approaches including bioinformatic models and analyses might enable deciphering of such complex signaling events, but require a system-wide molecular information at scale, which was previously out of reach for synaptic proteomes. This study bridges this gap and paves the way for the identification of key protein modules that underlie various synaptic functions. For example, by using different Cre-driver lines, the proteomes of various types of synapses can be obtained and correlated to any external synaptic features of interest, for example electrophysiological measurements. Similarly, we anticipate that multi-omic analysis of synapses will be conducted using an analogous experimental strategy, investigating the synaptic diversity of other biomolecules like glycans, lipids or RNA. Although transcriptomic analysis of FASS-sorted synaptosomes has been demonstrated only for the most abundant synapse-type (vGlut1+) (Hafner et al. 2019), RNA sequencing technology outperforms mass spectrometry in terms of absolute sensitivity and is predicted to accommodate synaptosome amounts lower than used in this study (Perez et al. 2021). However, the future integration of synaptic proteomes with local transcriptomes holds the promise to delineate the role of RNA localization in synaptic proteome diversity (Holt and Schuman 2013; Holt, Martin, and Schuman 2019). Here we provide a framework that can be developed to connect different levels of neurobiology, by combining synaptic proteome data with cellular or circuit-level experiments. On the cellular level, experiments are commonly targeted to a defined cell type or even synapse type to investigate function, connectivity or morphology using methods such as the Cre/lox system (Schroeder et al. 2023). With the strategy outlined here, such experiments can now be combined with synapse subtype specific proteome information and thereby connect different levels of organization. For example, one could probe how a particular phenotype, disease model, behavioral paradigm or cellular manipulation differentially affects the synaptic proteomes of the synapse subtypes of interest.

