## Supplementary Information for "The proteomic landscape of synaptic diversity across brain regions and cell types"

### Figure S1

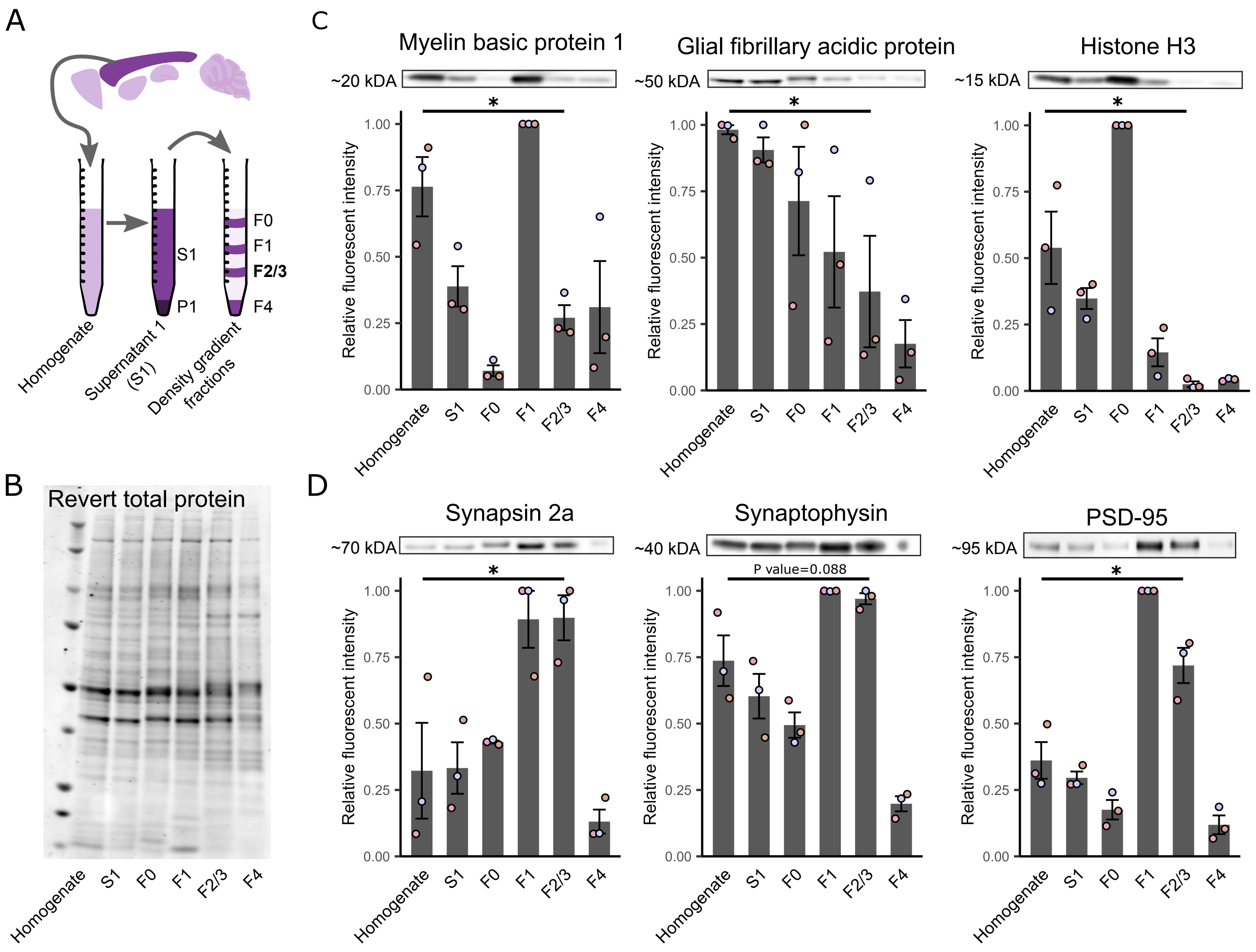

A
Homogenate
Supernatant 1 (S1)
Density gradient fractions
S1
P1
F0
F1
F2/3
F4
B
Revert total protein
Homogenate
S1
F0
F1
F2/3
F4
C
Myelin basic protein 1
~20 kDA
Glial fibrillary acidic protein
~50 kDA
Histone H3
~15 kDA
Relative fluorescent intensity
1.00
0.75
0.50
0.25
0.00
*
D
Synapsin 2a
~70 kDA
Synaptophysin
~40 kDA
P value=0.088
PSD-95
~95 kDA

A

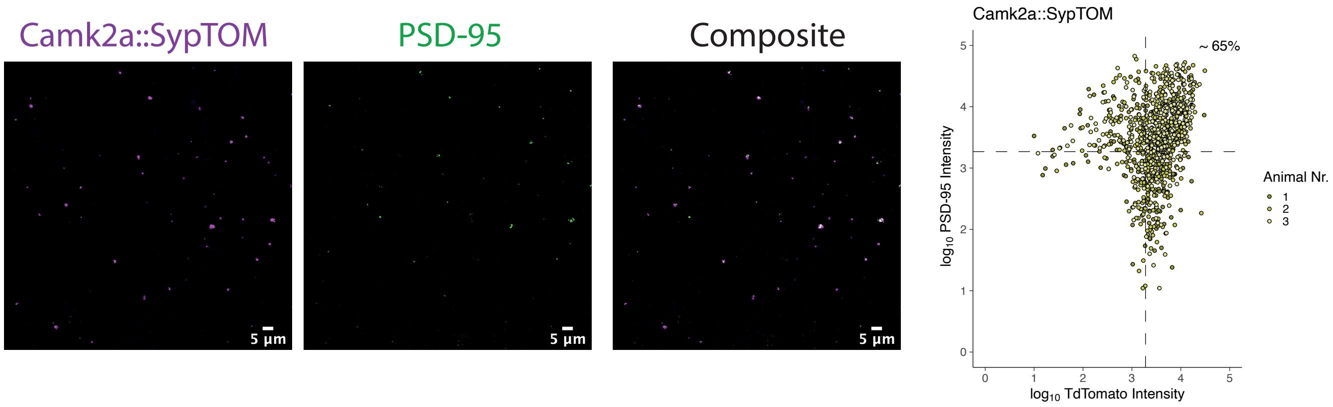

Camk2a::SypTOM
PSD-95
Composite
5 µm
5 µm
5 µm
Camk2a::SypTOM
~ 65%
log10 PSD-95 Intensity
log10 TdTomato Intensity
Animal Nr.
1
2
3


B

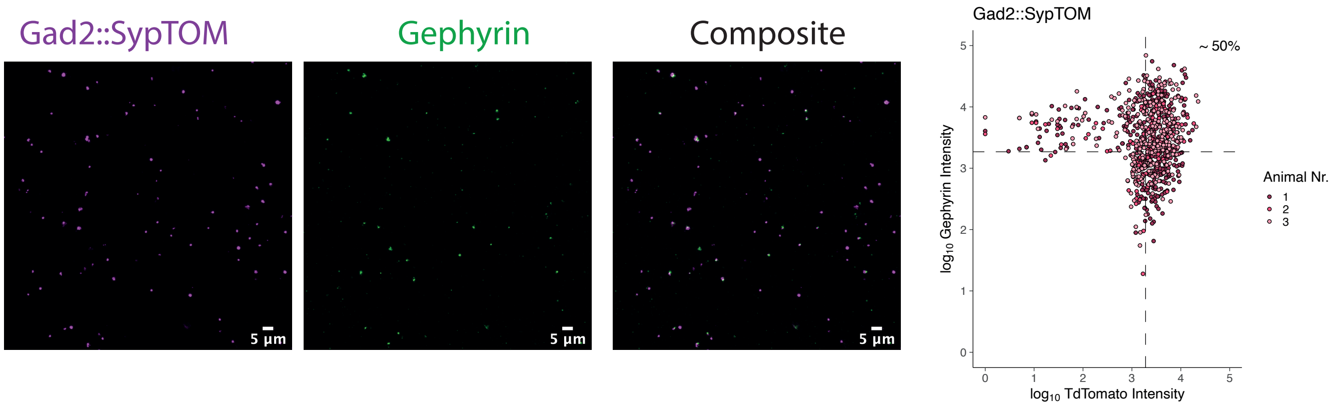

Gad2::SypTOM
Gephyrin
Composite
5 µm
5 µm
5 µm
Gad2::SypTOM
~ 50%
log10 Gephyrin Intensity
log10 TdTomato Intensity
Animal Nr.
1
2
3

### Figure S3.1

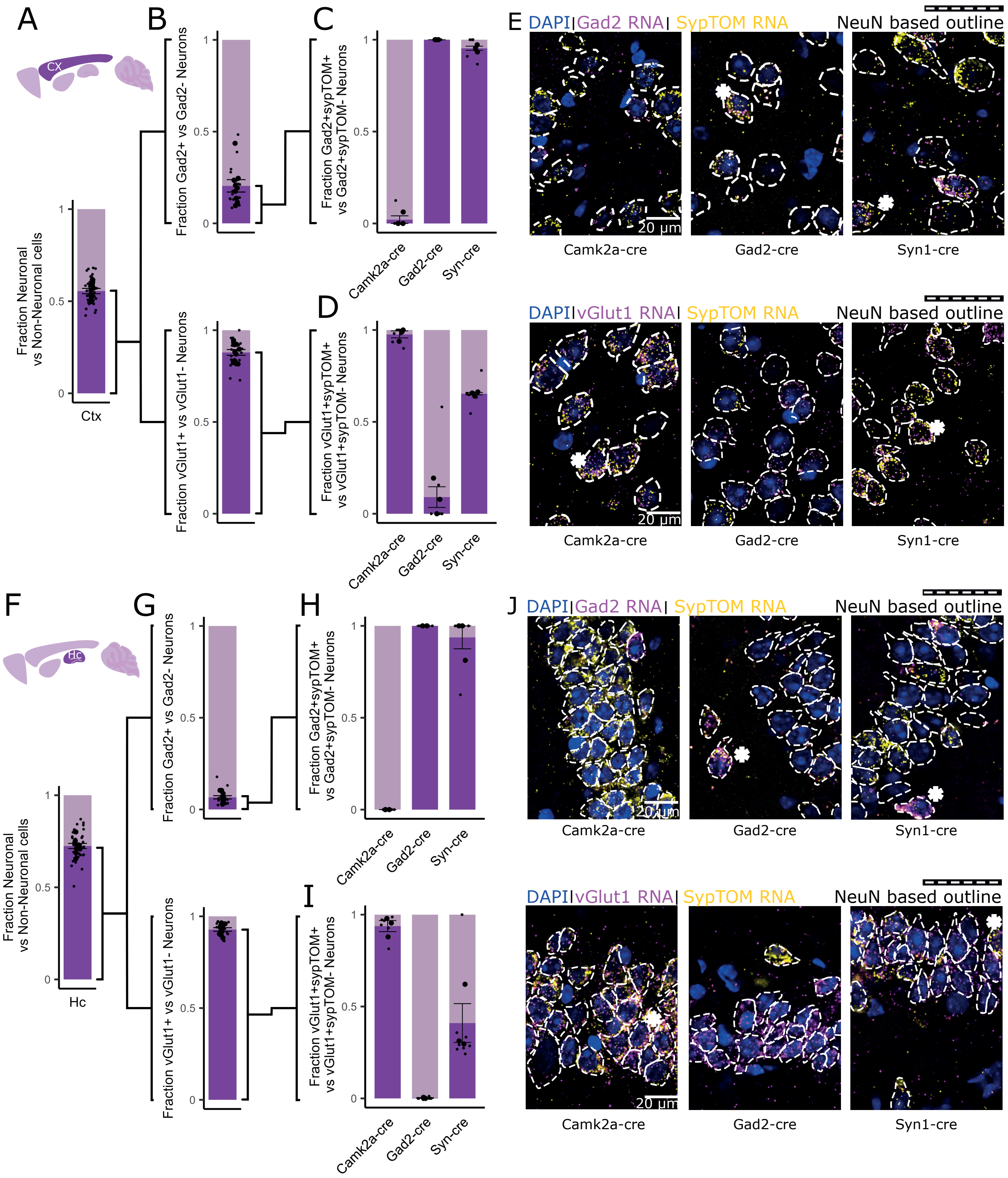

A
B
C
D
E
F
G
H
I
J
CX
Hc
Fraction Neuronal vs Non-Neuronal cells
Ctx
Hc
Fraction Gad2+ vs Gad2- Neurons
Fraction vGlut1+ vs vGlut1- Neurons
Fraction Gad2+sypTOM+ vs Gad2+sypTOM- Neurons
Fraction vGlut1+sypTOM+ vs vGlut1+sypTOM- Neurons
Camk2a-cre
Gad2-cre
Syn-cre
DAPI| Gad2 RNA| SypTOM RNA
DAPI| vGlut1 RNA| SypTOM RNA
NeuN based outline
20 µm
Camk2a-cre
Gad2-cre
Syn1-cre

### Figure S3.2

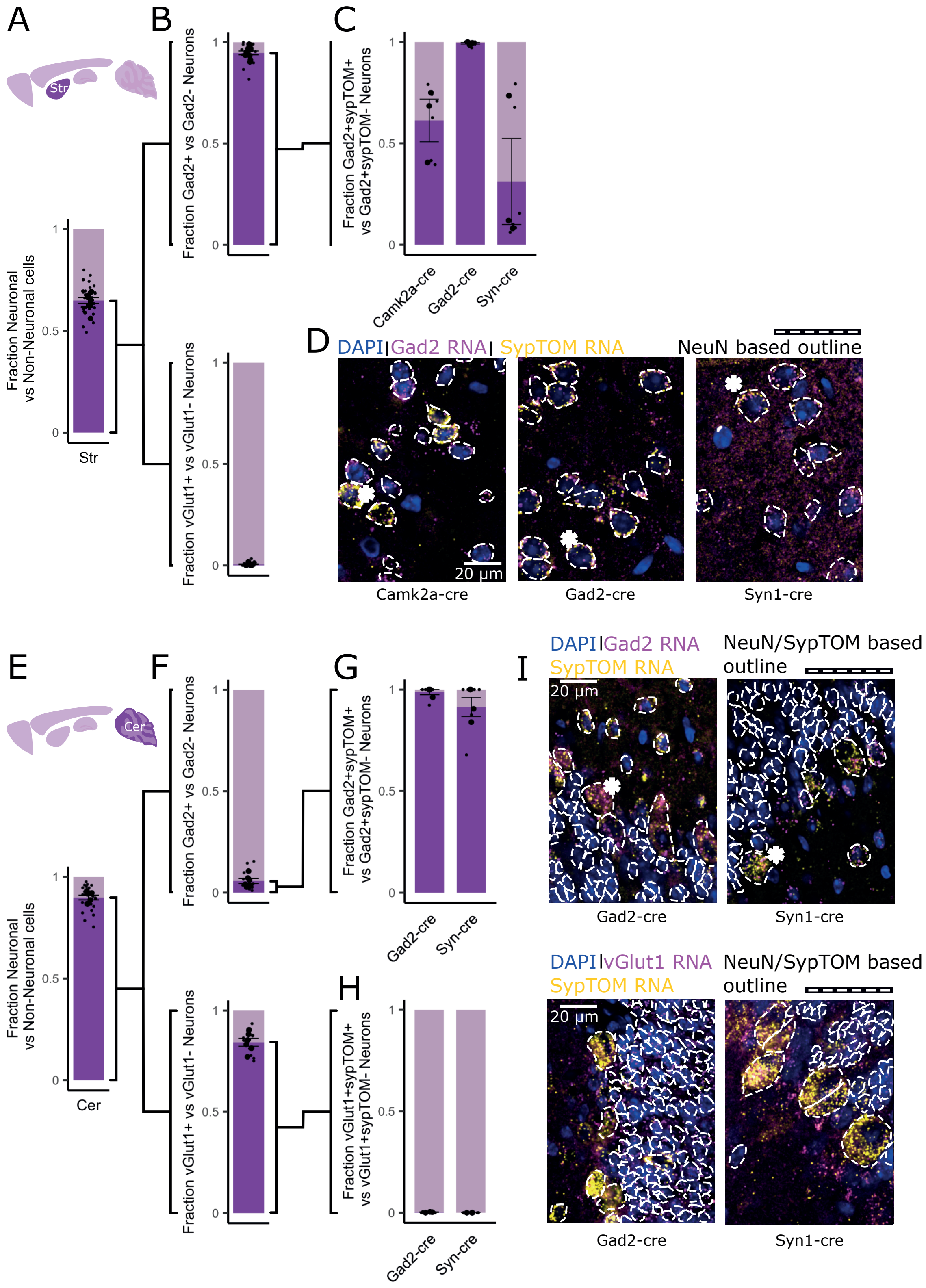

A
B
C
Str
Fraction Neuronal vs Non-Neuronal cells
Fraction Gad2+ vs Gad2- Neurons
Fraction Gad2+sypTOM+ vs Gad2+sypTOM- Neurons
Fraction vGlut1+ vs vGlut1- Neurons
Camk2a-cre
Gad2-cre
Syn-cre
D
DAPI | Gad2 RNA | SypTOM RNA
NeuN based outline
20 µm
Camk2a-cre
Gad2-cre
Syn1-cre
E
F
G
H
I
Cer
Fraction vGlut1+sypTOM+ vs vGlut1+sypTOM- Neurons
DAPI | Gad2 RNA
SypTOM RNA
NeuN/SypTOM based outline
DAPI | vGlut1 RNA
Syn1-cre

### Figure S3.3

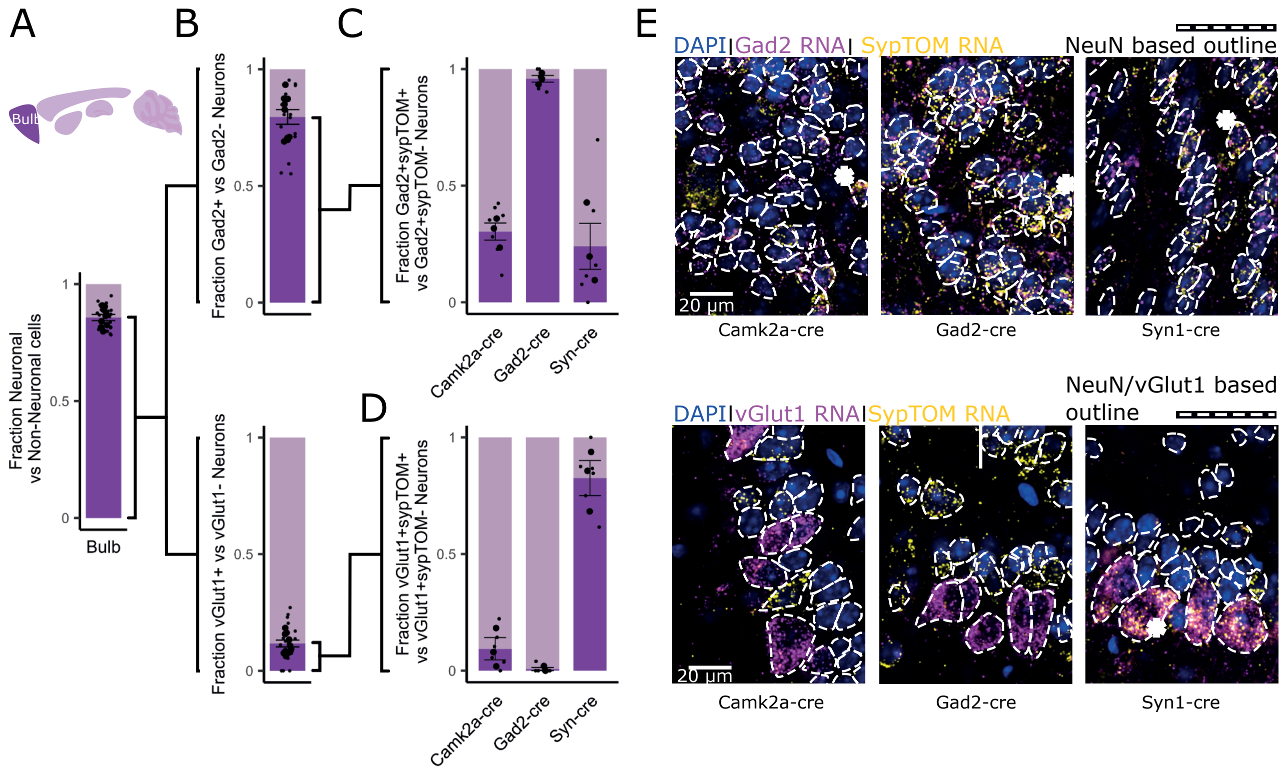
A
B
C
D
E
Bulb
Fraction Neuronal vs Non-Neuronal cells
Fraction Gad2+ vs Gad2- Neurons
Fraction Gad2+sypTOM+ vs Gad2+sypTOM- Neurons
Fraction vGlut1+ vs vGlut1- Neurons
Fraction vGlut1+sypTOM+ vs vGlut1+sypTOM- Neurons
Camk2a-cre
Gad2-cre
Syn-cre
DAPI|Gad2 RNA| SypTOM RNA
NeuN based outline
DAPI|vGlut1 RNA|SypTOM RNA
NeuN/vGlut1 based outline
20 µm
Syn1-cre

### Figure S4

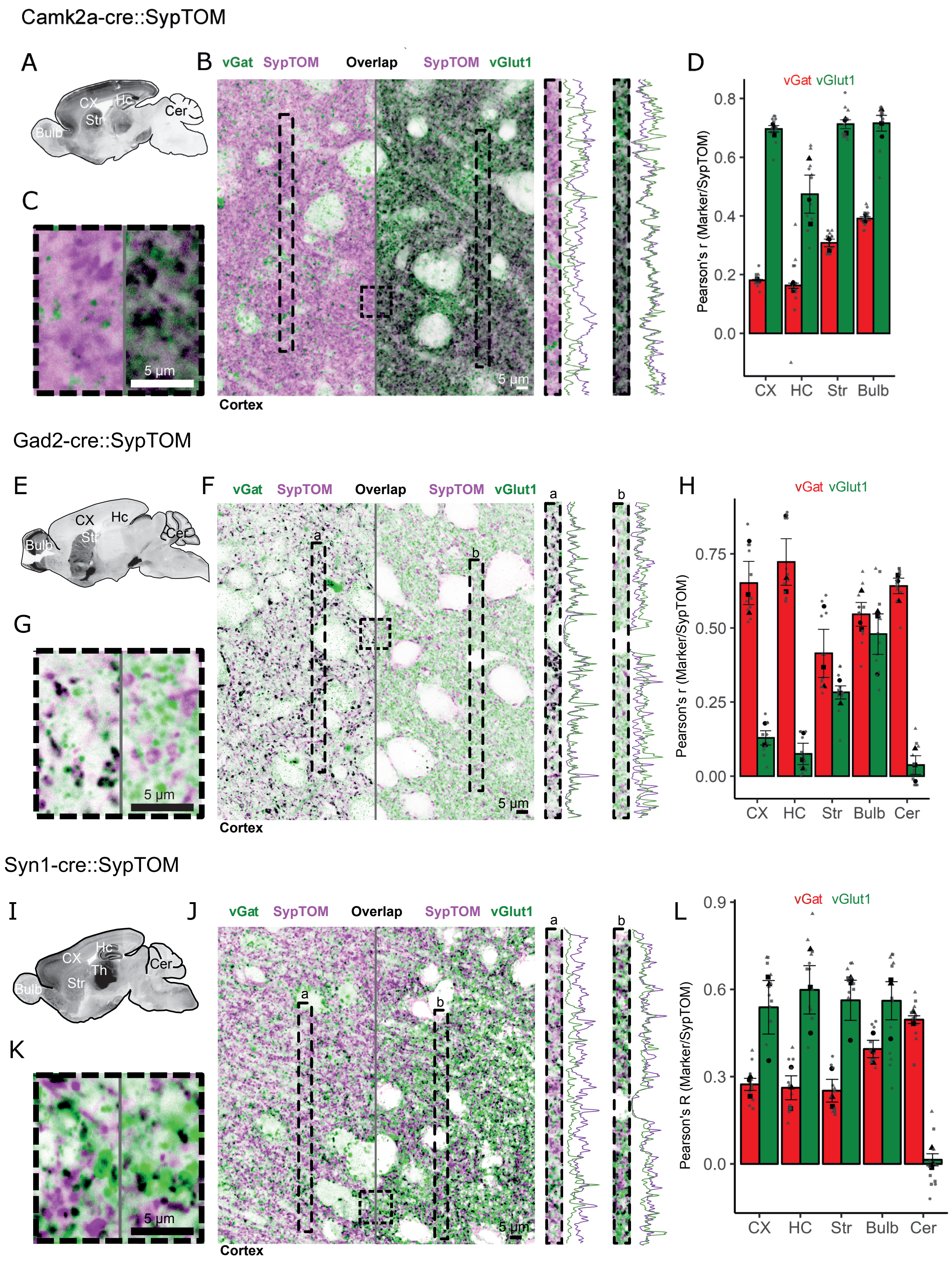

Camk2a-cre::SypTOM
A
CX
Hc
Str
Cer
Bulb
B
vGat SypTOM
Overlap
SypTOM vGlut1
5 µm
Cortex
C
5 µm
D
vGat vGlut1
Pearson's r (Marker/SypTOM)
0.8
0.6
0.4
0.2
0.0
CX HC Str Bulb
Gad2-cre::SypTOM
E
CX
Hc
Str
Cer
Bulb
F
vGat SypTOM
Overlap
SypTOM vGlut1
a
b
5 µm
Cortex
G
5 µm
H
vGat vGlut1
Pearson's r (Marker/SypTOM)
0.75
0.50
0.25
0.00
CX HC Str Bulb Cer
Syn1-cre::SypTOM
I
CX
Hc
Th
Str
Cer
Bulb
J
vGat SypTOM
Overlap
SypTOM vGlut1
a
b
5 µm
Cortex
K
5 µm
L
vGat vGlut1
Pearson's R (Marker/SypTOM)
0.9
0.6
0.3
0.0
CX HC Str Bulb Cer

### Figure S5

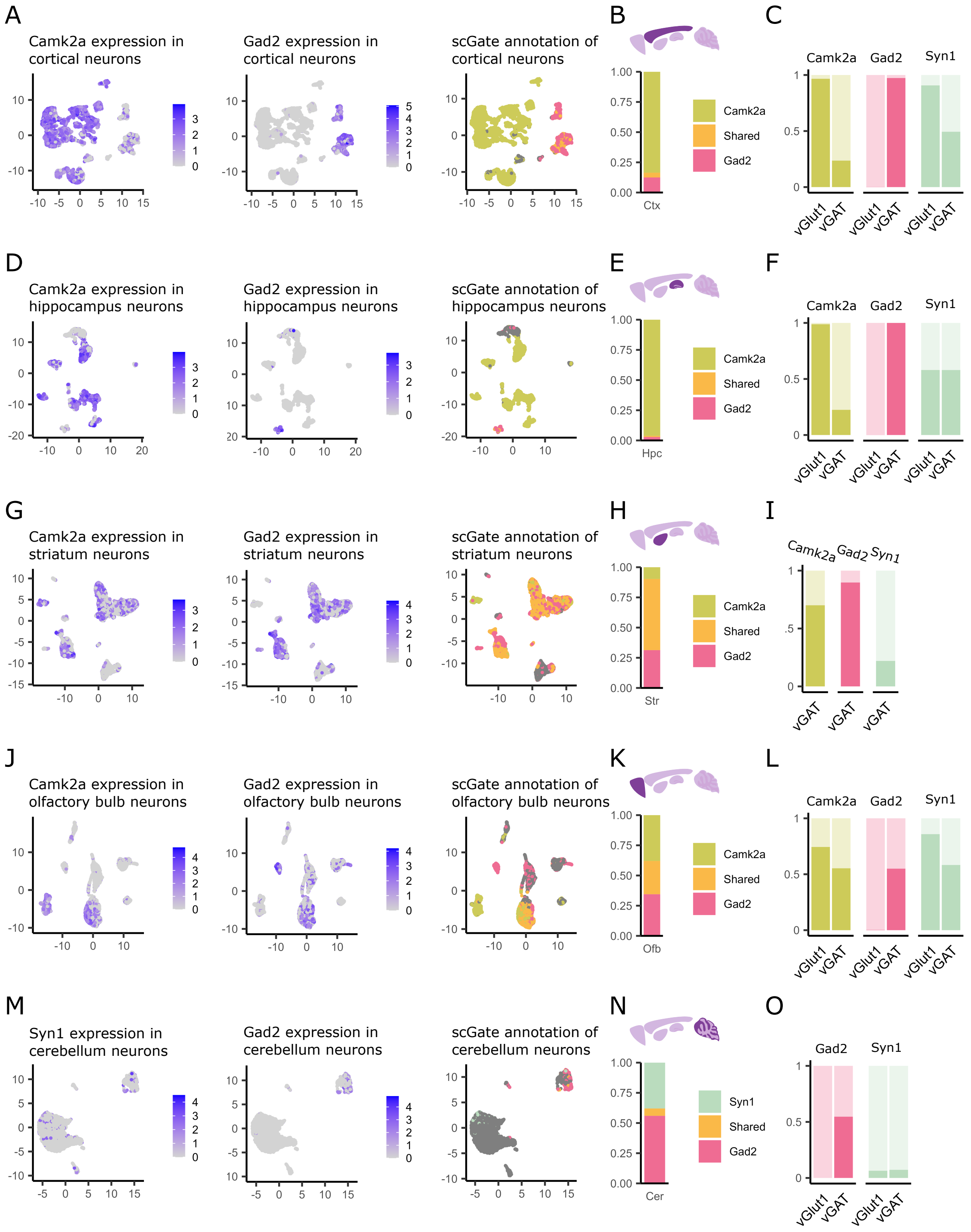

A
Camk2a expression in cortical neurons
Gad2 expression in cortical neurons
scGate annotation of cortical neurons
B
Ctx
Camk2a
Shared
Gad2
C
Camk2a
Gad2
Syn1
vGlut1
vGAT
D
Camk2a expression in hippocampus neurons
Gad2 expression in hippocampus neurons
scGate annotation of hippocampus neurons
E
Hpc
F
G
Camk2a expression in striatum neurons
Gad2 expression in striatum neurons
scGate annotation of striatum neurons
H
Str
I
J
Camk2a expression in olfactory bulb neurons
Gad2 expression in olfactory bulb neurons
scGate annotation of olfactory bulb neurons
K
Ofb
L
M
Syn1 expression in cerebellum neurons
Gad2 expression in cerebellum neurons
scGate annotation of cerebellum neurons
N
Cer
Syn1
O

### Figure S6

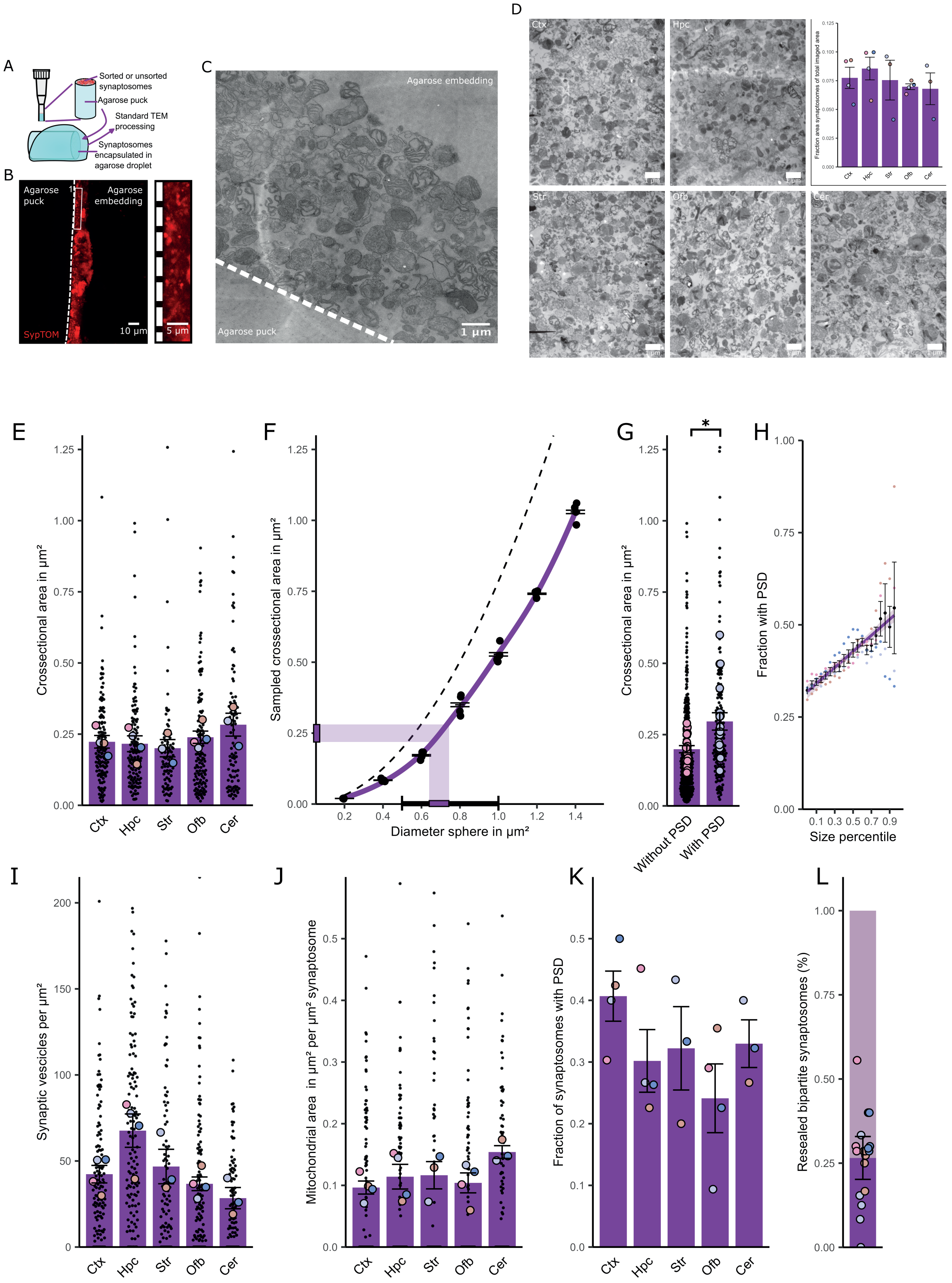

A
Sorted or unsorted synaptosomes
Agarose puck
Standard TEM processing
Synaptosomes encapsulated in agarose droplet
B
Agarose puck
Agarose embedding
SypTOM
10 µm
5 µm
C
Agarose embedding
Agarose puck
1 µm
D
Ctx
Hpc
Str
Ofb
Cer
E
Crossectional area in µm²
F
Sampled crossectional area in µm²
Diameter sphere in µm²
G
Crossectional area in µm²
Without PSD
With PSD
H
Fraction with PSD
Size percentile
I
Synaptic vescicles per µm²
J
Mitochondrial area in µm² per µm² synaptosome
K
Fraction of synaptosomes with PSD
L
Resealed bipartite synaptosomes (%)

Figure S7

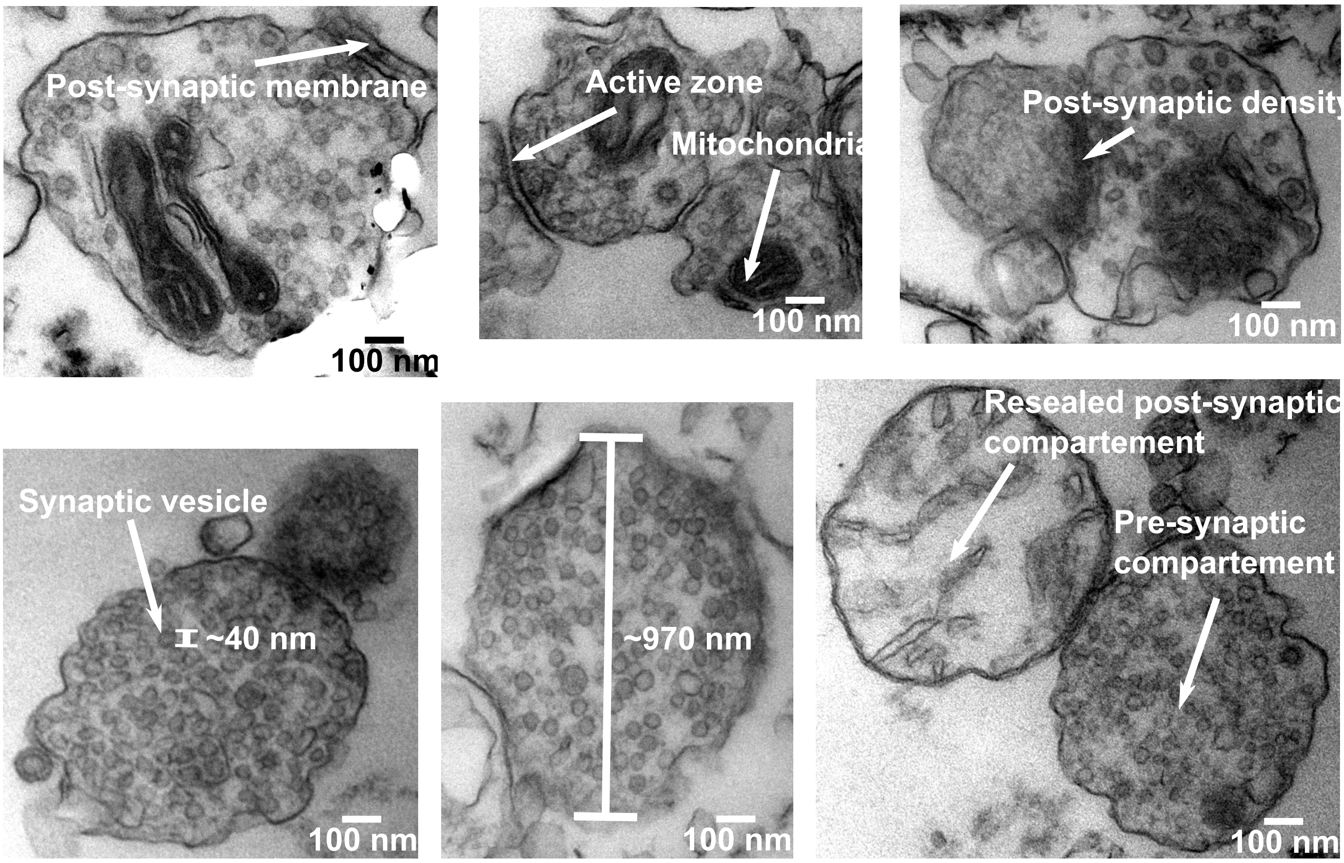

Post-synaptic membrane
100 nm
Active zone
Mitochondria
100 nm
Post-synaptic density
100 nm
Synaptic vesicle
~40 nm
100 nm
~970 nm
100 nm
Resealed post-synaptic compartement
Pre-synaptic compartement
100 nm

### Figure S8

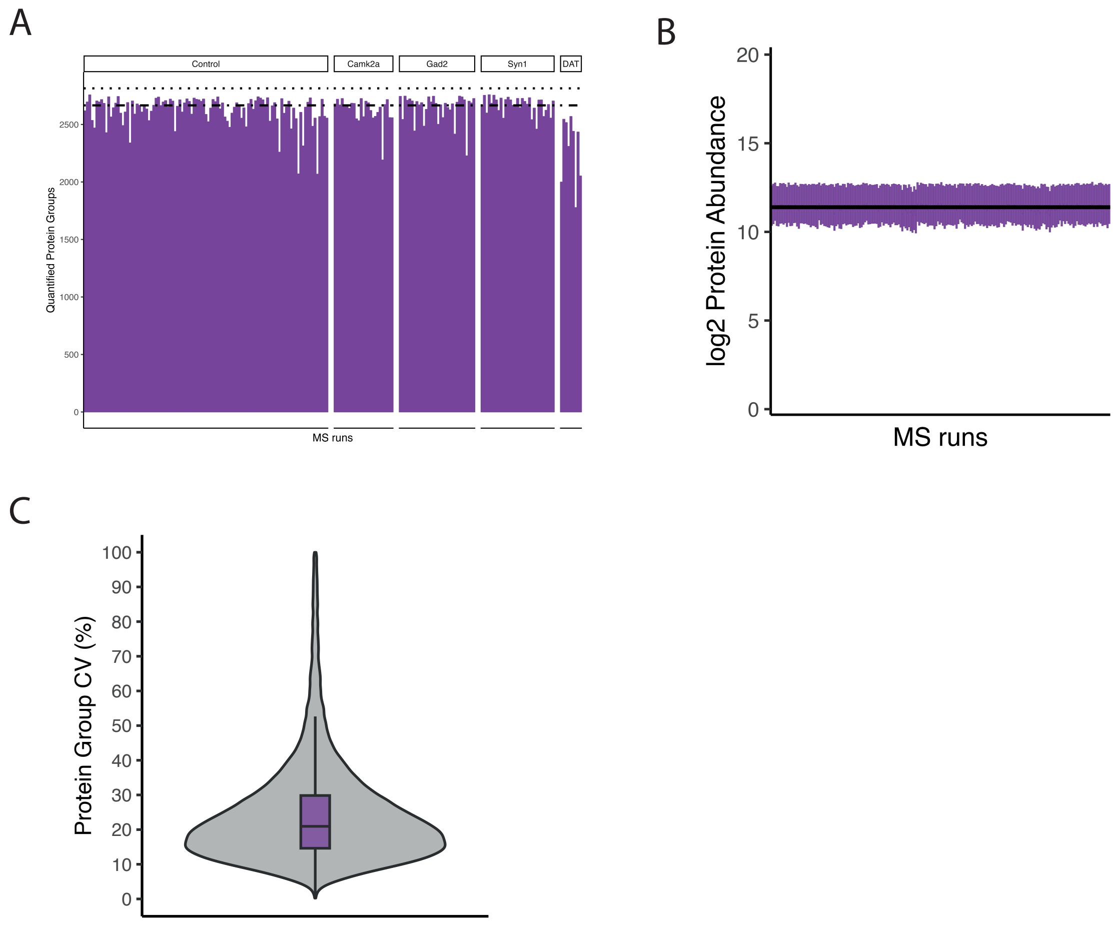

A
Control
Camk2a
Gad2
Syn1
DAT
Quantified Protein Groups
2500
2000
1500
1000
500
0
MS runs
B
log2 Protein Abundance
20
15
10
5
0
MS runs
C
Protein Group CV (%)
100
90
80
70
60
50
40
30
20
10
0

### Figure S9

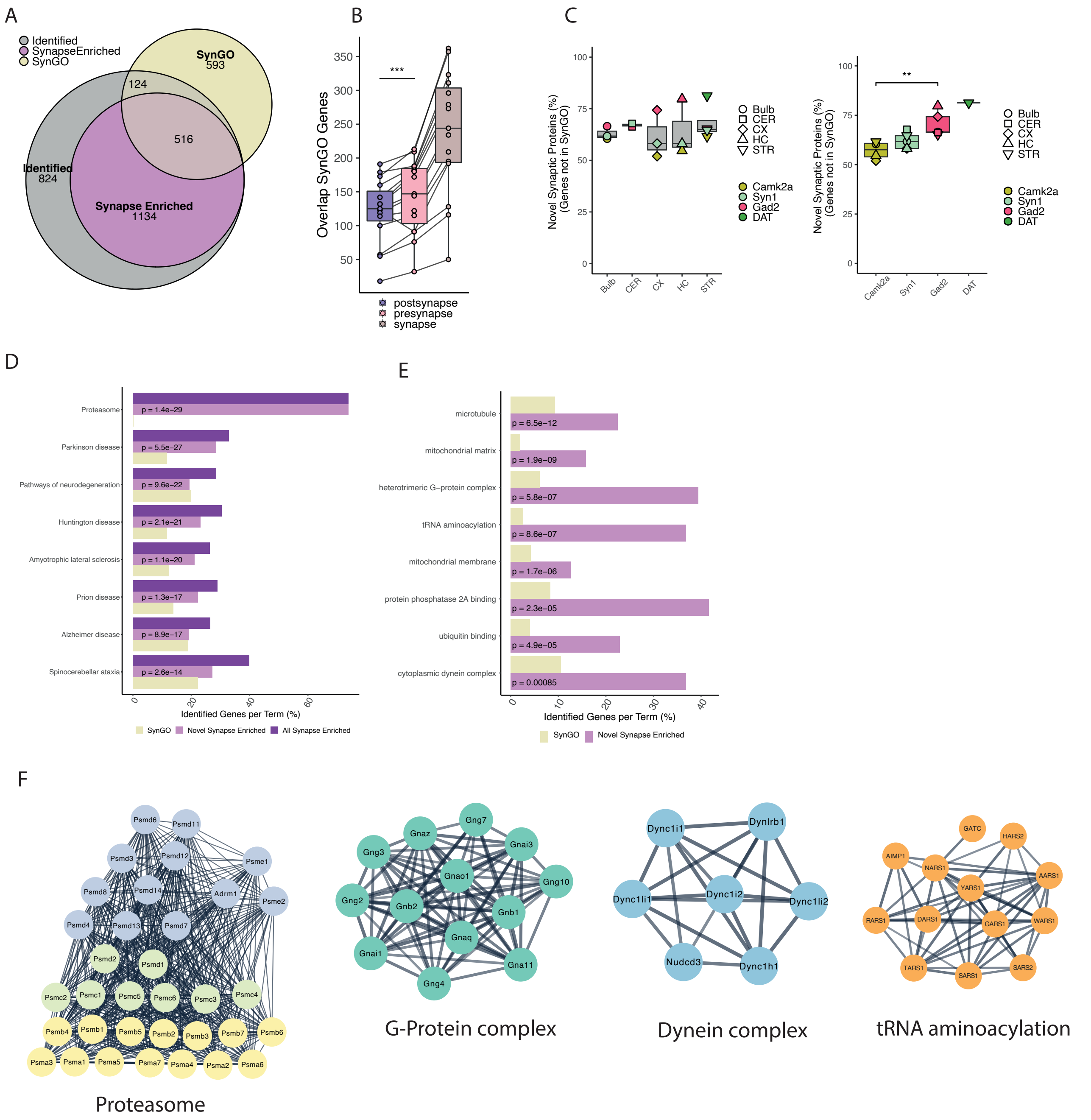

A
Identified
SynapseEnriched
SynGO
SynGO 593
124
516
Identified 824
Synapse Enriched 1134
B
Overlap SynGO Genes
***
postsynapse
presynapse
synapse
C
Novel Synaptic Proteins (%) (Genes not in SynGO)
Bulb
CER
CX
HC
STR
Camk2a
Syn1
Gad2
DAT
**
D
Proteasome
Parkinson disease
Pathways of neurodegeneration
Huntington disease
Amyotrophic lateral sclerosis
Prion disease
Alzheimer disease
Spinocerebellar ataxia
p = 1.4e−29
p = 5.5e−27
p = 9.6e−22
p = 2.1e−21
p = 1.1e−20
p = 1.3e−17
p = 8.9e−17
p = 2.6e−14
Identified Genes per Term (%)
SynGO
Novel Synapse Enriched
All Synapse Enriched
E
microtubule
mitochondrial matrix
heterotrimeric G−protein complex
tRNA aminoacylation
mitochondrial membrane
protein phosphatase 2A binding
ubiquitin binding
cytoplasmic dynein complex
p = 6.5e−12
p = 1.9e−09
p = 5.8e−07
p = 8.6e−07
p = 1.7e−06
p = 2.3e−05
p = 4.9e−05
p = 0.00085
Identified Genes per Term (%)
SynGO
Novel Synapse Enriched
F
Proteasome
G-Protein complex
Dynein complex
tRNA aminoacylation

### Figure S10

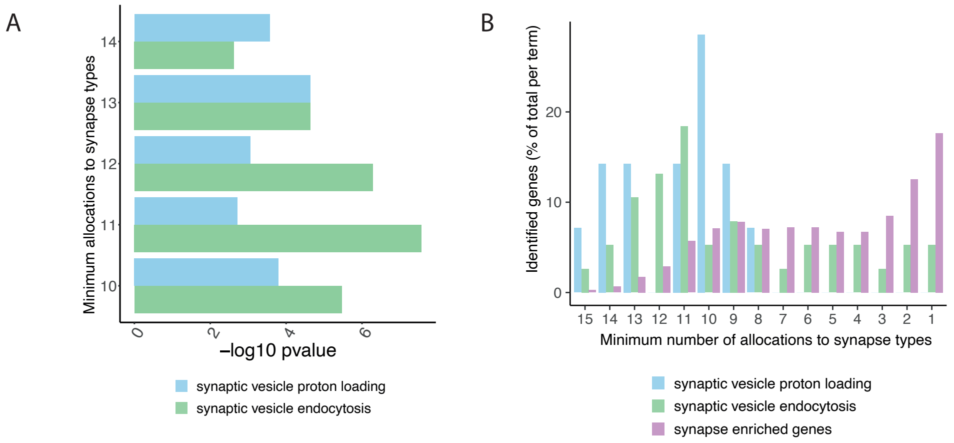
A
Minimum allocations to synapse types
14
13
12
11
10
0
2
4
6
−log10 pvalue
synaptic vesicle proton loading
synaptic vesicle endocytosis
B
Identified genes (% of total per term)
20
10
0
15 14 13 12 11 10 9 8 7 6 5 4 3 2 1
Minimum number of allocations to synapse types
synaptic vesicle proton loading
synaptic vesicle endocytosis
synapse enriched genes

### Figure S11

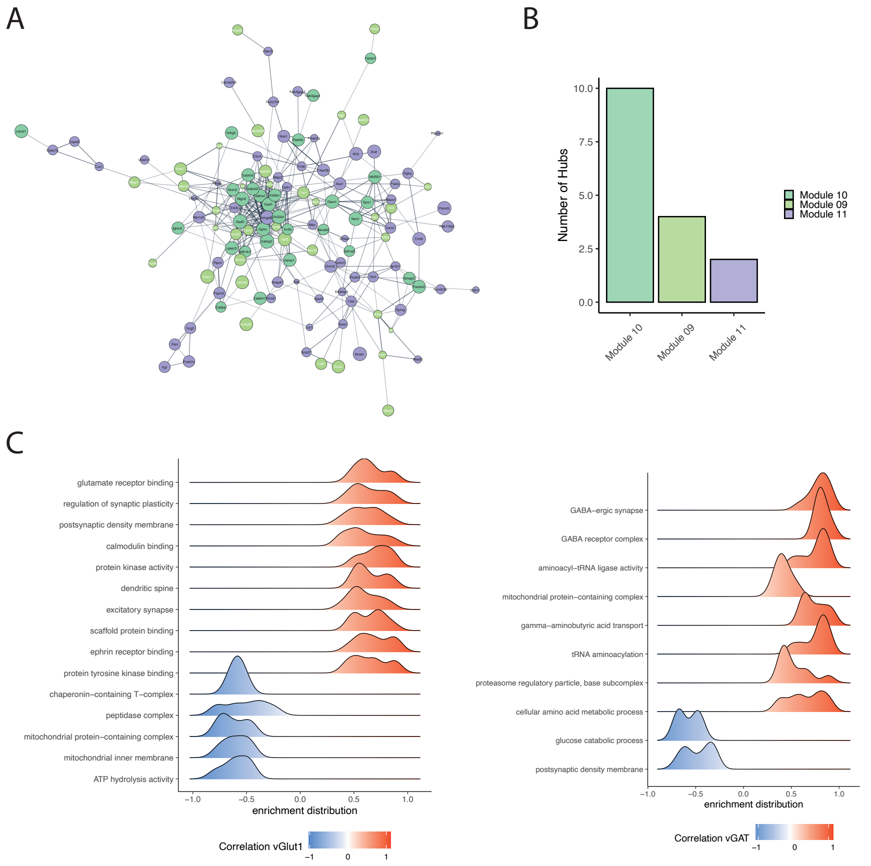

A
B
Number of Hubs
10.0
7.5
5.0
2.5
0.0
Module 10
Module 09
Module 11
C
glutamate receptor binding
regulation of synaptic plasticity
postsynaptic density membrane
calmodulin binding
protein kinase activity
dendritic spine
excitatory synapse
scaffold protein binding
ephrin receptor binding
protein tyrosine kinase binding
chaperonin−containing T−complex
peptidase complex
mitochondrial protein−containing complex
mitochondrial inner membrane
ATP hydrolysis activity
−1.0
−0.5
0.0
0.5
1.0
enrichment distribution
Correlation vGlut1
−1
0
1
GABA−ergic synapse
GABA receptor complex
aminoacyl−tRNA ligase activity
mitochondrial protein−containing complex
gamma−aminobutyric acid transport
tRNA aminoacylation
proteasome regulatory particle, base subcomplex
cellular amino acid metabolic process
glucose catabolic process
postsynaptic density membrane
enrichment distribution
Correlation vGAT

### Figure S12

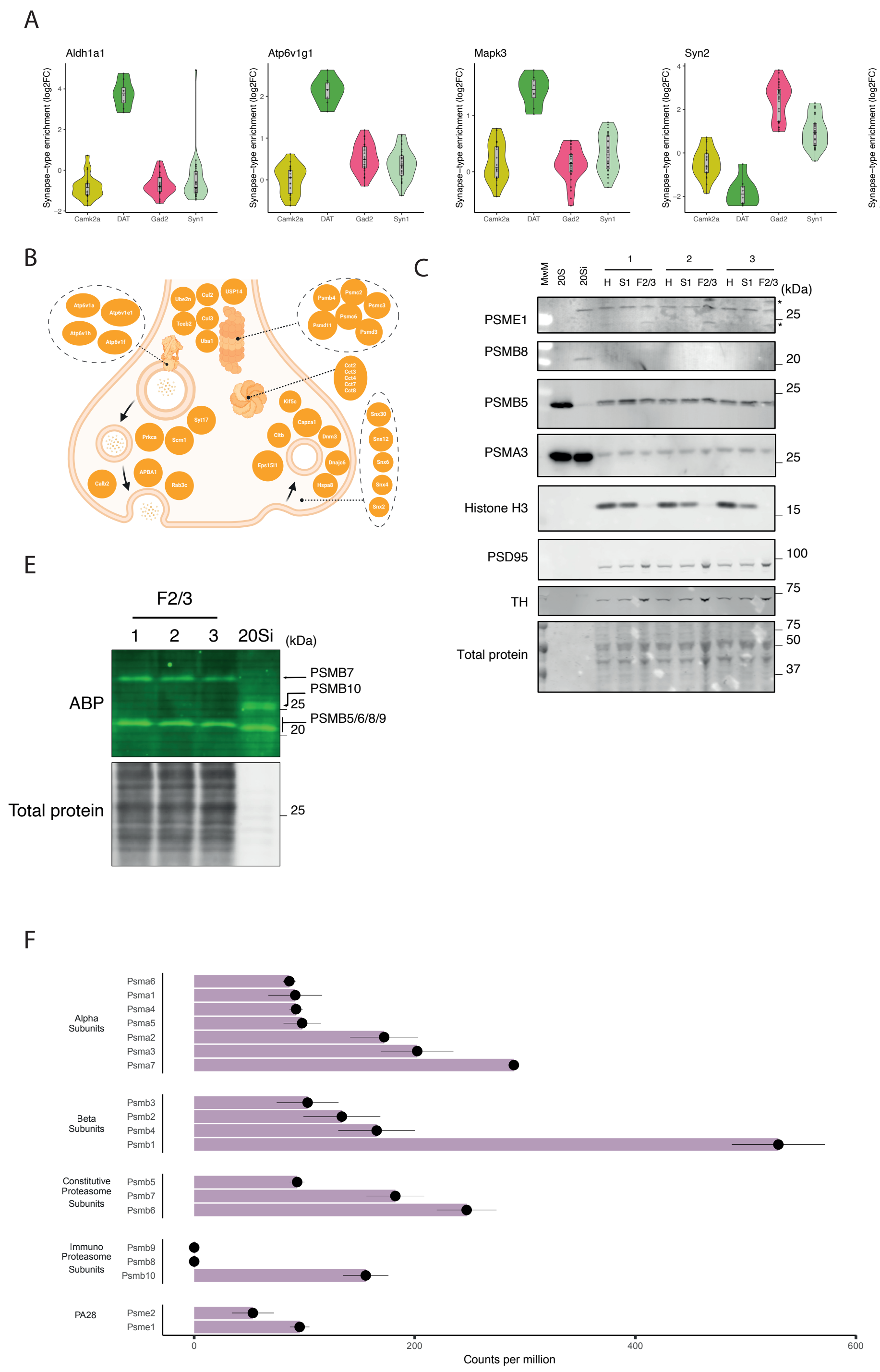
A
Aldh1a1
Atp6v1g1
Mapk3
Syn2
Psme1
Psme2
Synapse-type enrichment (log2FC)
Camk2a
DAT
Gad2
Syn1
B
C
PSME1
PSMB8
PSMB5
PSMA3
Histone H3
PSD95
TH
Total protein
MwM
20S
20Si
H
S1
F2/3
(kDa)
D
Silver staining
PSME1
PSMB5
Input
Beads
Isotype ctrl
PSME1 CST
PSME1 Abcam
E
ABP
Total protein
PSMB7
PSMB10
PSMB5/6/8/9
F
Alpha Subunits
Beta Subunits
Constitutive Proteasome Subunits
Immuno Proteasome Subunits
PA28
Counts per million

### Figure S13

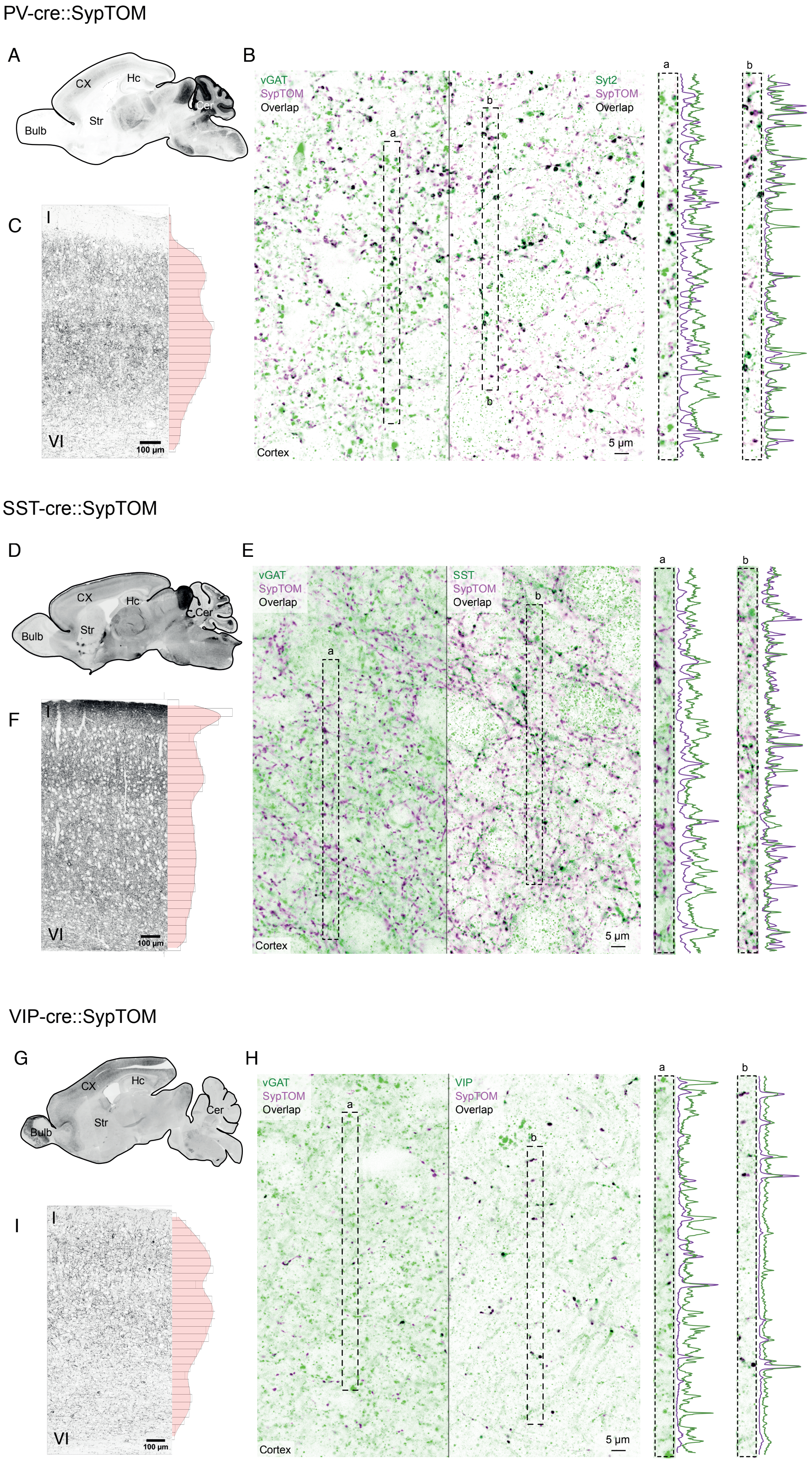

PV-cre::SypTOM
A
CX
Hc
Cer
Str
Bulb
B
vGAT
SypTOM
Overlap
Syt2
SypTOM
Overlap
a
b
5 µm
Cortex
C
I
VI
100 µm
SST-cre::SypTOM
D
CX
Hc
Cer
Str
Bulb
E
vGAT
SypTOM
Overlap
SST
SypTOM
Overlap
a
b
5 µm
Cortex
F
I
VI
100 µm
VIP-cre::SypTOM
G
CX
Hc
Cer
Str
Bulb
H
vGAT
SypTOM
Overlap
VIP
SypTOM
Overlap
a
b
5 µm
Cortex
I
I
VI
100 µm

### Figure S14

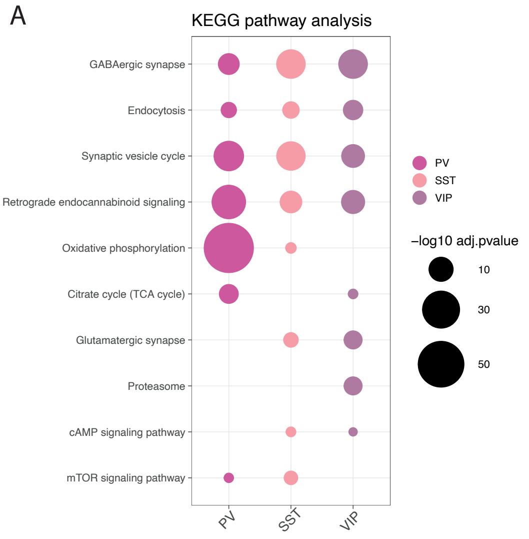

A
KEGG pathway analysis
GABAergic synapse
Endocytosis
Synaptic vesicle cycle
Retrograde endocannabinoid signaling
Oxidative phosphorylation
Citrate cycle (TCA cycle)
Glutamatergic synapse
Proteasome
cAMP signaling pathway
mTOR signaling pathway
PV
SST
VIP
−log10 adj.pvalue
10
30
50


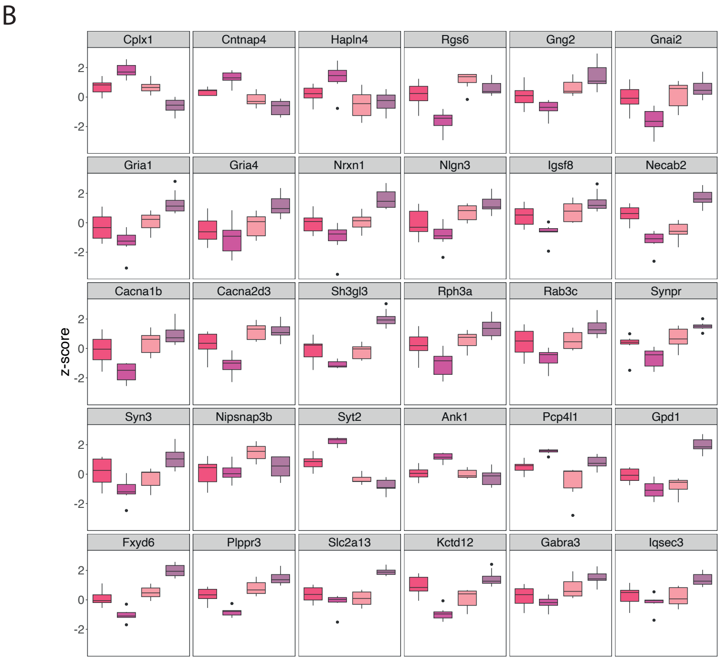

B
Cplx1
Cntnap4
Hapln4
Rgs6
Gng2
Gnai2
Gria1
Gria4
Nrxn1
Nlgn3
Igsf8
Necab2
Cacna1b
Cacna2d3
Sh3gl3
Rph3a
Rab3c
Synpr
Syn3
Nipsnap3b
Syt2
Ank1
Pcp4l1
Gpd1
Fxyd6
Plppr3
Slc2a13
Kctd12
Gabra3
Iqsec3
z-score

***Supplementary Figure legends***

**Figure S1. SDS-PAGE and Immunoblot of synaptosome preparation reveals enrichment of synaptic proteins and depletion of non-synaptic contaminants.**

(A) Schematic depiction of the Percoll gradient synaptosome preparation. Homogenates of murine cortex were centrifuged resulting in a pellet 1 (P1) and supernatant 1 (S1), which was subsequently layered onto the Percoll gradient and centrifuged yielding different fractions. F2/3 represents the synaptosomal fraction that occurs at the interface between 23% and 10% Percoll.

(B) Representative revert™ 700 total protein stain used to normalize for differences in protein loading.

(C) Contaminant proteins were depleted in the synaptosomal fraction; Bar plots depict the relative fluorescent intensity across the different fractions of Myelin basic protein 1 (Paired one tailed t-test homogenate vs F2/3, $p = 0.023$; n=3), the glial marker glial fibrillary acidic protein (Paired one tailed t-test homogenate vs F2/3, $p = 0.048$; n=3) and the nuclear marker Histone H3 (Paired one tailed t-test homogenate vs F2/3, $p = 0.032$; n=3).

(D) Synaptic proteins were enriched in the synaptosomal fraction; synapsin 2a (Paired one tailed t-test homogenate vs F2/3, $p = 0.024$; n=3), synaptophysin (Paired one tailed t-test homogenate vs F2/3, $p = 0.088$; n=3) and postsynaptic density protein 95 (Paired one tailed t-test homogenate vs F2/3, $p = 0.017$; n=3). Representative immunoblot bands are shown above the corresponding bars. Error bars represent the standard error of the mean. Relative fluorescent intensity is normalized for protein loading, local fluorescent background and the maximum intensity of each sample.

**Figure S2. Immunofluorescence of spotted synaptosomes shows the percentage of synaptosomes that include a postsynaptic element.**

(A) Camk2a+ synaptosomes were spotted onto gelatin-covered coverslips after FASS and immunostained for the PSD95 protein. The left panel includes representative images of spotted synaptosomes and their TdTomato fluorescence and PSD95 immunoreactivity. The right graph depicts a scatterplot showing PSD95 and TdTomato fluorescent intensity for each identified spot. Percentage of double-positive fraction is indicated on the top left. n=3 animals.

(B) The same as in A but for Gad2+ synaptosomes and Gephyrin immunoreactivity.

**Figure S3.1. Characterization of cell types labeled by Camk2a-cre::SypTOM, Gad2-cre::SypTOM and Syn1-cre::SypTOM mouse lines in cortex and hippocampus by RNA fluorescent *in situ* hybridisation (FISH) on sagittal brain slices.**

Camk2a-cre::SypTOM and Gad2-cre::SypTOM specifically label vGlut1+ and Gad2+ neurons respectively, while Syn1-cre labels both types of neurons, in the cortex and hippocampus.

(A) Depicts the fractions of neurons, as defined by RNA binding fox-1 homolog 3 (NeuN), vs non-neuronal cells, as defined by NeuN negative and DAPI+, in the cortex. Big data points represent individual mice (n=9), while small data points represent individual images (4 per mouse).

(B) The top graph depicts the fraction of neurons expressing Gad2 (n=9, 18 images), and the bottom graph the fraction of neurons expressing vGlut1 (n=9, 18 images).

(C) Shown is the fraction of Gad2+ neurons labeled by the different Cre driver lines (n=3, 6 images for each driver); Note the minimal labeling by the Camk2a-cre driver line.

(D) Displayed is the fraction vGlut+ neurons that were labeled by the different cre-driver lines (n=3, 6 images for each driver); Note the minimal labeling by Gad2-cre and the incomplete labeling by Syn1-cre.

(E) Representative single molecule FISH images where the dashed line represents the outline of the NeuN signal, yellow dots the SypTOM RNA, purple dots the respective markers RNA and in blue nuclei stained with DAPI. Asterisks signify examples of co-expression, error bars the standard error of the mean.

(FGHIJ) The same as ABCDE but for the hippocampus.

**Figure S3.2. Characterization of cell types labeled by Camk2a-cre::SypTOM, Gad2-cre::SypTOM and Syn1-cre::SypTOM mouse lines in striatum and cerebellum by RNA fluorescent *in situ* hybridization (FISH) on sagittal brain slices.**

(A) Depicts the fraction of neurons, as defined by RNA binding fox-1 homolog 3 (NeuN) or morphology (for horizontal neurons of the olfactory bulb and purkinje neurons in the cerebellum, which are known to lack NeuN expression). Shown is the fraction of neurons vs non-neuronal cells in the striatum. Large data points represent individual mice (n=9) and smaller points depict individual images (4 per mouse).

(B) The top graph shows that the vast majority of striatum neurons expressed Gad2 RNA, while the bottom graph shows that vGlut1+ neurons were virtually absent.

(C) Shown is the fraction of Gad2+ neurons labeled by the different cre-drivers. All drivers labeled Gad2+ neurons but to varying degrees; while Gad2-cre labels all Gad2+ neurons, Camk2a-cre and Syn1-cre only labeled subpopulations.

(D) Representative single molecule FISH images where the dashed line represents the outline of the NeuN signal, the yellow dots the SypTOM RNA, the purple dots the respective markers RNA and in blue nuclei stained with DAPI. Asterisks signify examples of co-expression, error bars the standard error of the mean (SEM).

(EFGHI) Similar to ABCD above. vGlut1+ neurons make up the majority of neurons in cerebellum. These were not labeled by Gad2-cre and Syn1-cre while Gad2+ neurons were labeled almost completely.

Note that most excitatory synapses labeled in striatum (by Camk2a-cre and Syn1-cre) and cerebellum (by Syn1-cre) originate in other brain regions because excitatory neuronal cell bodies are either not present (striatum) or not labeled (cerebellum).

**Figure S3.3. Characterization of cell types labeled by Camk2a-cre::SypTOM, Gad2-cre::SypTOM and Syn1-cre::SypTOM mouse lines in the olfactory bulb by RNA fluorescent *in situ* hybridisation (FISH) on sagittal brain slices.**

(A) Depicts the fraction of neurons, as defined by RNA binding fox-1 homolog 3 (NeuN) or morphology (for horizontal neurons of the olfactory bulb and purkinje neurons in the cerebellum, which are known to lack NeuN expression). Shown is the fraction of neurons vs non-neuronal cells in the olfactory bulb. Large data points represent individual mice (n=9) and smaller points depict individual images (4 per mouse).

(B) The top graph depicts the fraction of neurons expressing Gad2, and the bottom graph the fraction of neurons expressing vGlut1.

(C) Shown is the fraction of Gad2+ neurons labeled by the different cre-driver.

(D) Representative single molecule FISH images where the dashed line represents the outline of the NeuN signal, the yellow dots the SypTOM RNA, the purple dots the respective markers RNA and in blue nuclei stained with DAPI. Asterisks signify examples of co-expression, error bars the standard error of the mean. Note that Camk2a-cre, Syn1-cre and Gad2-cre label Gad2+ neurons in the olfactory bulb, although, Camk2a-cre and Syn1-cre less so. vGlut+ neurons are lowly abundant in the olfactory bulb and are only strongly labeled by Syn1-cre.

**Figure S4. Characterization of the synapses that are labeled by Camk2a-cre::SypTOM, Gad2-cre::SypTOM and Syn1-cre::SypTOM mouse lines by assessment of overlap with vGat and vGlut1 immunoreactivity in sagittal brain slices.**

Excitatory synapses are fluorescently labeled with high specificity in the isocortex and hippocampus in Camk2a-cre::SypTOM mice.

(A) Sagittal overview of SypTOM expression in a Camk2a-cre::SypTOM mouse; Fluorescent signal (black) was strong in the cortex (Ctx), hippocampus (Hc), striatum (Str) and less strong in the olfactory bulb (Ofb).

(B) Representative image of a immunofluorescent co-staining for the inhibitory synapse marker Solute Carrier Family 32 Member 1 (vGat) and the excitatory synapse marker Solute Carrier Family 17 Member 7 (vGlut1), both in green, and of SypTOM expression, in purple, in the Ctx of a Camk2a-cre::condSypTOM mouse. Overlap is depicted in black; SypTOM overlapped mainly with vGlut1, which is further illustrated by maximum normalized fluorescent intensity line plots (a,b).

(C) Magnification of the staining depicted in B.

(D) Pearson's correlation between SypTOM fluorescence and the respective markers show distinct labeling in the Ctx and Hc (n=3 animals, 4 images per brain region for every marker and mouse), and to a lesser extent in Str and Ofb. Larger data points represent a mouse, while smaller points depict individual images. Error bars signify the standard error of the mean (SEM).

(EFGH) The same analysis as in ABCD but for the Gad2-cre::SypTOM mouse, and additionally for the cerebellum. The Gad2-cre::SypTOM mouse shows specific labeling for inhibitory synapses in cortex, hippocampus and cerebellum and lower specificity in the striatum and olfactory bulb.

(IJKL) The same analysis as in EFGH but for Syn1-cre::SypTOM. The Syn1-cre::SypTOM mouse shows overlapping synaptic labeling with both vGat and vGlut1 in all brain regions except for the cerebellum, where predominantly inhibitory synapses are labeled.

**Figure S5. Re-analysis of published scRNA sequencing data corroborates the characterization of cell types labeled in the different Cre-driver line crosses by RNA FISH.**

(A) UMAP plots of cortical neurons from Zeisel et. al. (Zeisel et al. 2018) colored for normalized Camk2a and Gad2 RNA expression in the left and middle panel respectively. The right panel for both using annotation by scGate (Andreatta, Berenstein, and Carmona 2022).

(B) Quantification of the neurons annotated as Camk2a+, Gad2+ or Camk2a+ & Gad2+ shows that Camk2a+ and Gad2+ neurons are predominantly distinct populations.

(C) Co-expression of excitatory marker Solute Carrier Family 17 Member 7 (vGlut1) and inhibitory marker Solute Carrier Family 32 Member 1 (vGat) demonstrates that Camk2a is expressed in most vGlut1+ neurons, while Gad2 is expressed in most vGat+ neurons.

(DEF) Hippocampal neurons are similar to cortical neurons in these regards; Camk2a and Gad2 expression is predominantly distinct and expressed in most vGlut1+ and vGat+ neurons respectively.

(GHI) In contrast, Camk2a and Gad2 expressions overlap substantially in striatal neurons and are both expressed in vGat+ neurons, while the number of vGlut1 expressing neurons is minimal (not shown).

(JKL) In olfactory bulb neurons there is overlap between Camk2a+ and Gad2+ populations. Camk2a expression is present in both vGlut1+ and vGat+ neurons, while Gad2 expression is confined to vGat+ neurons.

(MNO) In the cerebellum Gad2 expression is confined to the vGat+ neurons, while Syn1 expression is low in both vGat+ and vGlut+ neurons.

**Figure S6. Synaptosomes from different brain areas do not differ in their ultrastructural features.**

(A) Schematic representation of the processing of synaptosomes for electron microscopy (EM). Synaptosomes are spun down onto an agarose puck lodged in a pipette tip and, subsequently, encapsulated in agarose.

(B) Confocal images of synaptosomes purified by FASS, spun down onto the agarose puck and embedded in additional agarose.

(C) Unsorted synaptosomes are structurally preserved during the processing; Displayed is an electron micrograph tile scan showing synaptosomes, and other particles, on the agarose puck. The edge of the puck is depicted by the dashed line.

(D) Synaptosome density in electron micrographs is comparable across the analyzed brain regions. Displayed are representative EM tile scans (acquired with a 31500X magnification) of the synaptosome fraction originating from the cortex (Ctx), hippocampus (Hpc), Striatum (Str), olfactory bulb (Ofb) and cerebellum (Cer). Scale bars represent one micrometer. Synaptosomes are identifiable as round structures containing synaptic vesicles. The bar chart shows the fractional area of the image which could be annotated as synaptosome. We find no significant differences across the analyzed brain regions (repeated measures ANOVA, $p = 0.95$; Ctx: n=4, Hpc: n=4; Str: n=3; Ofb: n=4; Cer: n=4; originating from 4 animals).

(E) Synaptosomes from different brain areas have the same size. Cross sectional area of synaptosomes in $\mu m^2$ originating from cortex (Ctx), hippocampus (Hpc), Striatum (Str), olfactory bulb (Ofb) and cerebellum (Cer) are not significantly different (Repeated measures ANOVA, $p = 0.21$; Ctx: n=4, Hpc: n=4; Str: n=3; Ofb: n=4; Cer: n=4; originating from 4 animals). Larger data points represent the average cross-sectional area from 30 synaptosomes, while smaller data points depict individual synaptosomes

(F) Estimation of synaptosome size assuming that synaptosomes are spheres. Synaptosome cross-sectional sampling results in an underrepresentation of the true size. Therefore, we estimated the extent of underrepresentation of the true synaptosomes size assuming spherical shape.We randomly sampled 30 planes from a sphere with the indicated diameter (n=4) and plotted the obtained average cross-sectional area, represented by the black dots in the plot. Connecting the sampled data points provides the purple line which was used to estimate the synaptosome diameter from the experimentally sampled mean cross-sectional area. The dashed line indicates the maximal cross sectional area per diameter assuming sphericity without cross-sectional sampling. The horizontal bar on the X-axis indicates the size range previously reported for synaptosomes (0.5 to 1 $\mu m^2$) (Gulyássy et al. 2020). The average cross sectional area for all brain regions (light purple area) predict a synaptosome size within the established range for synaptosomes.

(G) Synaptosomes that have a postsynaptic density (PSD) have a higher cross-sectional area, indicating that their cross-sectional sampling is closer to the maximum (Paired one tailed t-test synaptosomes with PSD vs without PSD, $p = 0.031$; n=4; brain regions pooled).
(H) Conversely, the fraction of synaptosomes with a PSD attached scales with cross-sectional synaptosome area percentile; Plotted is the fraction of synaptosomes with a PSD for different percentiles. A linear line is fitted with shading indicating 99% confidence interval.
(I) The number of synaptic vesicles, normalized for synaptosome size, did not differ significantly across the analyzed brain regions (Repeated measures ANOVA, $p = 0.083$; same samples as in A).
(J) Mitochondrial area, normalized for synaptosome size, was not significantly different between brain regions (Repeated measures ANOVA, $p = 0.337$; same samples as in A).
(K) The fraction of synaptosomes with a visible PSD in the cross-section was not significantly different between brain regions (Repeated measures ANOVA, $p = 0.15$; same samples as in A).
(L) Bipartite synaptosomes (synaptosomes containing both a pre- and postsynaptic element) have a resealed postsynaptic compartment in ~25% of the cases; Colors of the data points indicate that the synaptosomes originate from the same mice, albeit different brain regions. All error bars indicate standard error of the mean.

**Figure S7. Synaptosome integrity is preserved during fluorescence-activated synaptosome sorting.**
Representative images of sorted synaptosomes acquired at a magnification of 4100X. Recognizable ultrastructural features are annotated as well as the size of a synaptic vesicle (~40nm) and the diameter of a synaptosome (~970 nm). Scale Bar represents 100 nm.

**Figure S8. Mass-spectrometry based proteomic analysis of 15 synapse types.**
(A) Barplot showing the number of quantified protein groups per sample, sorted by cell types. Dash-dotted line indicates the mean number of quantified protein groups and dotted line indicates the total number of quantified proteins in the experiment.
(B) Log2 protein intensities across all MS samples. Black dots indicate median intensity, upper and lower hinges the 25th and 75th percentile.
(C) Violin plot showing the distribution of protein group coefficients of variation (CV) within conditions (synapse types), the median CV is ~20%. Boxplot indicates the median, 25th and 75th percentile.

**Figure S9. Comparison of Synapse-enriched proteins with the SynGO database.**

(A) Euler diagram showing overlap between proteins that were quantified in the experiment, proteins that were identified as synapse-enriched and the SynGO database. 81% of the SynGO annotated proteins that were identified in the experiment were identified as synapse-enriched.

(B) Boxplot showing overlap with SynGO-annotated genes for all synapse types and the terms “synapse”, “presynapse” and “postsynapse”. Each dot represents a synapse subtype. Overlap with SynGO was significantly higher for the term “presynapse” compared to the term “postsynapse”. n=15, paired t-test, ***P<0.001.

(C) Boxplots showing percentage of novel synaptic proteins stratified by brain region (left) or by cell type (right). We detected significantly more novel synaptic proteins at Gad2+ synapses compared to Camk2a+ synapses. n=4 (Camk2a), 5 (Gad2), t-test, **P<0.01.

(D) Barplot showing selected significantly enriched KEGG pathways for all genes in SynGO, all synapse-enriched proteins and the proteins identified as synapse enriched but not previously annotated in SynGO. X-axis shows the percentage of identified genes per term. Text indicates p-value of the “novel synapse enriched” group.

(E) The same as in D but for selected Gene Ontology terms.

(F) Visualization of selected proteins that are associated with enriched terms in the “novel synapse enriched” group using data from the STRING interaction database. Edges represent stringdb score >0.7 (high confidence).

**Figure S10. Synaptic vesicle endocytosis and vATPases are enriched among the proteins that are found at the most synapse types.**

(A) Barplot showing SynGO enrichment p-values for the proteins that were identified at minimally 10, 11, 12, 13 or 14 of the total of 15 synapse types.

(B) Distribution of proteins that were associated with the SynGO terms “synaptic vesicle proton loading” or “synaptic vesicle endocytosis” depending on groups defined by the minimum number of synapse types they are identified in. The distribution for all synapse enriched protein is plotted for comparison.

**Figure S11. Validation and further characterization of the protein-protein correlation network.**

(A) STRING network of proteins that are associated with the three modules that are significantly correlated with vGat (module 9,10,11). Edges are based on stringdb score >0.4.

(B) Barplot showing number of hubs from the network in A for each module. Hubs are defined as the top 10% most connected proteins in the network (centrality parameter “degree”). The core module (module 10) of the protein-protein correlation network also contains the most hubs in the STRING network from A.

(C) Ridgeplots for pathways enriched in vGlut1 and vGat protein communities. Ridgeplots show enrichment distribution for core enriched genes of selected significantly enriched terms. Gene set enrichment analysis (GSEA) was conducted using Gene Ontology terms and proteins ranked by their correlation with vGat (right) and vGlut1 (left) immunofluorescence.

**Figure S12. Supplementary analysis of the dopaminergic synaptic proteome**

(A) Violin plots for two representative proteins showing specific enrichment or specific depletion in dopaminergic synaptic terminals compared to all other synapse types.

(B) Scheme showing selected proteins that were enriched in Dat+ synapses as well as Syn1+ synapses within the presynaptic terminal.

(C) SDS-PAGE and immunoblot of the different fractions (H: homogenate, S1: soluble fraction, F2/3: synaptosomal fraction) of the synaptosome preparations from mouse striata (n=3 biological replicates). While in the F2/3 fractions synaptic proteins PSD95 and TH were enriched, the nuclear protein Histone H3 was not detectable. The constitutive subunits of the proteasome PSMA3 and PSMB5 as well as the PA28 regulatory particle subunit PSME1 were found across all fractions at comparable levels. By contrast, the immunoproteasome subunit PSMB8 was only be detected in the purified 20Si proteasome sample.

(D) Immunoblot of a PSME1 co-immunoprecipitation experiment on primary rat cortical neurons. The experiment was performed with two different antibodies against PSME1 along with an isotype control and naked beads. The experiment shows that in rat cortical neurons PSME1 interacts with the constitutive 20S proteasome.

(E) SDS-PAGE of mouse striatal F2/3 fractions and purified 20Si proteasome assayed for proteasome activity using an activity-based probe (ABP). While activity corresponding to the PSMB9 subunit of the 20Si proteasome is visible in the purified sample, this is not the case in the F2/3 striatal fractions. The asterisks indicate non-specific bands.

(F) Re-analysis of published scRNA sequencing data (Zeisel et al. 2018) showing expression of proteasomal genes in midbrain dopaminergic neurons. mRNA for the constitute proteasome subunits are detected while mRNA of 2 of 3 immunoproteasome subunits are not detected.

**Figure S13. Characterization of the synapses that are labeled by PV-cre::SypTOM, SST-cre::SypTOM and VIP-cre::SypTOM mouse lines in sagittal brain slices.**

(A) Sagittal overview of SypTOM expression in a PV-cre::SypTOM mouse.

(B) Representative image of a immunofluorescent co-staining for the inhibitory synapse marker Solute Carrier Family 32 Member 1 (vGat) and the synaptotagmin-2 (Syt2), a marker for synapses formed by PV-neurons (Sommeijer and Levelt 2012), both in green, and of SypTOM expression, in purple, in the Ctx

of a PV-cre::condSypTOM mouse. Overlap is depicted in black; SypTOM overlaps with Syt2 and partially with vGat, which is further illustrated by maximum normalized fluorescent intensity line plots (a,b).
(C) Overview of SypTOM expression in the cortex of a PV-cre::SypTOM mouse. The density across the different cortical layers is plotted on the side.
(DEF) Same as ABC but for the SST-cre::SypTOM mouse and the SST-neuron marker protein somatostatin.
(GHI) Same as ABC but for the VIP-cre::SypTOM mouse and the VIP-neuron marker protein VIP peptides.

**Figure S14. Supplementary analysis for cortical interneuron subtype proteomes.**
(A) KEGG pathway enrichment analysis of cortical interneuron proteomes.
(B) Boxplots for representative proteins that show specific enrichment in the indicated cortical interneuron subtypes.

**Acknowledgements**
We thank the MPI Brain Research Imaging Facility, Florian Vollrath, Christine Molenda, Ángeles Macias Pardo and Stephan Junek for assistance with microscopy and preparation of samples for electron microscopy. We thank all members of the MPI Brain Research Proteomics Facility for maintenance and assistance with data acquisition. We thank all members of the MPI Brain Research Animal Facility and Guido Schmalbach and Fabian Bayer from the MPI Brain Research workshop. We are grateful to the members of the Schuman lab research group for discussion and support at all stages of the project. We thank Ina Bartnik, Teresa Spano and Belquis Nassim Assir for technical support with experiments; Etienne Herzog and his research group for sharing a Fiji macro to analyze synaptosome immunofluorescence; and Julia Kuhl for graphical support. For funding, we acknowledge the Swiss National Science Foundation SNSF (Postdoc Mobility fellowships P2EZP3_191820 and P400PB_199288 for M.v.O.), the Max Planck Society and the European Union (ERC, DiverseSynapse, 101054512). Views and opinions expressed are however those of the author(s) only and do not necessarily reflect those of the European Union or the European Research Council. Neither the European Union nor the granting authority can be held responsible for them.

**Author contributions**
M.v.O. performed all experiments except those noted below and co-wrote the manuscript. T.B. performed immunoblotting, histology, electron microscopy experiments and reanalysis of scRNA sequencing data. S.L.G. performed Proteasome validation experiments. S.t.D. contributed to methodology and analysis. G.T.

contributed to data analysis. N.F. performed synaptosome preparations. J.L. supervised proteomics. E.M.S. supervised the project and co-wrote the manuscript. All authors edited the manuscript.

#### Material and Methods

##### Animals

All animal procedures were executed in accordance with institutional guidelines and approved by the relevant authorities. The following Cre-driver lines were crossed with the SypTOM (JAX strain #: 012570, Ai34D) mouse line: Camk2a-cre (Tsien et al. 1996) (JAX strain #: 005359), Gad2-cre (Taniguchi et al. 2011) (JAX strain #: 010802), Syn1-cre (Zhu et al. 2001) (JAX strain #: 003966), Dat-cre (Bäckman et al. 2006) (JAX strain #: 006660), PV-cre (Hippenmeyer et al. 2005) (JAX strain #: 008069), SST-cre (Taniguchi et al. 2011) (JAX strain #: 013044) and VIP-cre (Taniguchi et al. 2011) (JAX strain #: 010908). Animals were housed under a 12 h light/dark cycle, and provided with food and water ad libitum.

##### Histology

Mice were transcardially perfused with 4% paraformaldehyde (PFA) in PBS and brains were post-fixed overnight in 4% PFA at 4 °C. Sagittal sections (50µm) were cut using a Leica vibratome (VT1200S) and washed in PBS. Vibratome sections were blocked and permeabilized by incubation in 5% goat serum 0.5% Triton-x in PBS (blocking buffer) at RT for 4 hours. Primary antibody incubation was performed ON at 4 °C on a rocker in blocking buffer. The following day, sections were washed in PBS and incubated with a fluorescently-labeled secondary antibody ON at 4 °C. Slices were washed in PBS, rinsed in ddH2O and air dried on SuperFrost Plus glass slides (Fisher Scientific). Aqua-Poly/Mount (Polysciences) was used for mounting.

For single molecule fluorescent *in-situ* hybridization experiments (viewRNA, Thermo Fisher), mice were perfused and postfixed with 4% PFA, 4% sucrose in PBS for 1 hour at room temperature (RT) instead. Cryoprotection was performed by incubating ON in 15% and subsequently 30% sucrose in RNAse free PBS at 4 °C. Sagittal sections (40µm) were cut using a Zeiss HYRAX S50 (at -25 °C) and washed in PBS. Sections were post-fixed at RT in a 4% PFA solution (4% paraformaldehyde, 5.4% Glucose, 0.01 M sodium metaperiodate, in lysine-phosphate buffer) for 10 minutes, washed in RNAse free PBS and permeabilized using a detergent solution (viewRNA) for 20 minutes at RT. Probe hybridization was performed in the hybridization buffer at 40 °C ON. The following day, sections were rinsed in the wash buffer and subjected sequentially to pre-amp DNA, amp DNA and label probe oligonucleotides in their respective buffers for 1 hour at 40 °C with washes in the wash buffer at RT in between. After

washing in PBS, sections were blocked in blocking buffer (4% goat serum 0.5% Triton-X in PBS). Primary antibody staining was performed ON in blocking buffer at 4 °C. After washing in PBS, secondary antibody incubation was carried out at RT for 2 hours in blocking buffer. Sections were washed, counterstained with DAPI (1 µg/ml, Thermo Fisher) in PBS for 3 minutes, washed in PBS and mounted as described above. The following primary antibodies and corresponding dilution factors were used: anti-vGAT (SYSY, 131004 – gp, 1:5000), anti-vGAT (SYSY, 131002, rb, 1:5000), anti-vGlut1 (SYSY, 135304, gp, 1:5000), anti-vGlut1 (SYSY, 135303, rb, 1:5000), anti-TH (SYSY, 213104, gp 1:500), anti-VIP (Thermo Fisher, PA5-78224, rb, 1:500), anti-SST (Thermo Fisher, PA5-85759, rb, 1:500), anti-Syt2 (SYSY, 105223, rb, 1:500), anti-NeuN (Abcam, Ab177487, rb, 1:1000). The following FISH probes were used at 1:100 dilution: vGlut1 (viewRNA, VB1-15833-VC, Alexa Fluor 647), Gad2 (viewRNA, VB6-17621-VC, Alexa Fluor 750), TdTomato (viewRNA, VF1-14985, Alexa Fluor 647), TdTomato (viewRNA, VF6-13925, Alexa Fluor 750).
Images were acquired using a Zeiss confocal microscope (LSM-880 or LSM-980) a 63X or 40X objective (NA 1.4).

Synaptosome preparation

Synaptosomes were prepared as described in Westmark et. al. (Westmark et al. 2011). Briefly, animals were sacrificed by decapitation, the brain regions of interest were dissected on ice and subsequently homogenized in gradient medium (GM; 0.25 M sucrose, 5mM Tris-HCl, 0.1mM EDTA supplemented with Calbiochem Protease inhibitor cocktail III) using a glass dounce homogenizer. The homogenate was centrifuged for ten minutes at 4 °C at 1’000g. The supernatant (S1) was layered onto a Percoll density gradient with 23%, 10% and 3% Percoll in GM buffer. The gradient was centrifuged for 5min at 32’500g at 4 °C with maximum acceleration and minimum deceleration using a Beckman Coulter JA-25.50 rotor in an Avanti J-26S XPI centrifuge (both from Beckman Coulter). The resulting bands are labeled, from top to bottom, F0, F1, F2/3, F4. Bands were retrieved and directly processed (for EM analysis) or stored at -20 °C. For proteasome activity measurement, F2/3 fractions were filtered onto glass fiber filters using a syringe and stored at -80 °C until further processing. For immunoblotting, synaptosomes were lysed in a detergent buffer (8M urea, 10% SDS, 10% Sodium deoxycholate 5% Triton-X in water; 1 part detergent buffer and 5 parts synaptosome fraction) at 75 °C for 5 minutes. BCA assay (Thermo Fisher) was used to approximate protein concentrations of the different fractions.

Proteasome activity assay

After fraction collection on filters proteasome activity was assayed by incubation of filters with HR buffer (50 mM Tris–HCl pH 7.4, 5 mM MgCl2, 250 mM sucrose) (de Jong et al. 2012) freshly supplemented with 1 mM DTT, 2 mM ATP and 1 µM Me4BodipyFL-Ahx3Leu3VS (Bio-Techne GmbH, I-190-050) for 1h at 37 C. To assay proteasome activity of 20S (LifeSensors, PS020) and 20Si (Enzo, BML-PW9645-0050) purified samples, 1 µg of protein was incubated in HR buffer complete with 1 µM Me4BodipyFL-Ahx3Leu3VS for 1h at 37 °C. Sample protein concentration was measured with the precision red advanced protein assay (Cytoskeleton, Inc., ADV02-A).

SDS-PAGE and immunoblotting

For each sample, a volume corresponding normalized protein amounts was supplemented with 10X SDS sample buffer (500 mM Tris pH 6.8, 25% SDS and 2% bromophenol blue in 70% glycerol-30% dH2O), NuPAGE Sample Reducing Agent (10X) (Thermo Fisher, NP0004) and distilled water to an equal final volume. Samples were denatured and reduced at 90 °C for 5 minutes and run on Novex Tris-Glycine 4-20% and Novex 12% Bis-Tris mini gels. Activity-based probe fluorescence was measured either in gel or after transfer on the membranes on an Azure Sapphire biomolecular imager. For silver staining the gels were processed as described in the kit's technical bulletin (Thermo Fisher, 24612). For immunoblot analyses, the gels were wet-transferred onto Immobilon-FL PVDF membranes (Sigma-Aldrich, 05317-10EA). Equal loading and transfer were then assessed by Revert 700 total protein stain (LI-COR, 926-11011) or Ponceau stain (CST, 59803). The membranes were destained, blocked for 1h at room temperature in Intercept (TBS) blocking buffer (LI-COR, 927-60001) and probed with primary antibodies overnight at 4 °C. The next day the membranes were developed with fluorescently-labeled secondary antibodies on a LI-COR Odyssey Classic system. The following primary antibodies and corresponding dilution factors were used: anti-PSMA3 (Enzo, BML-PW8110-0100 – ms, 1:1000), anti-PSMB8 (Enzo, BML-PW8845-0100 – ms, 1:1000), anti-PSMB5 (CST, 12919 – rb, 1:1000), anti-PSME1 (Abcam, ab155091 – rb, 1:1000), anti-Histone H3 (Abcam, ab1791 – rb, 1:1000), anti-TH (SYSY, 213104 – gp, 1:1000), anti-PSD95 (Abcam, ab2723 – ms, 1:1000, used for Proteasome validation), anti-PSD95 (Thermo Fisher, MA1-046, ms, 1:1000, used for synaptosome prep analysis), anti-Mbp (Abcam, Ab62631– ms, 1:1000), anti-GFAP (abcam, Ab7260 - rb, 1:1000), anti-Syn (SYSY, 106002, rb, 1:1000), anti-SYPH (Sigma, S5768, ms, 1:5000).

Electron microscopy

The procedure for electron microscopy (EM) analysis of unsorted synaptosomes was based on Sebring et. al. (Sebring, Johnson, and Spall 1988) with chemicals from Sigma Aldrich and Plano GmbH (Pioloform). F2/3 fractions were fixed with EM grade glutaraldehyde (final concentration 2.5%) for 30 minutes on ice and, subsequently, diluted in PBS. During the fixation period, 5% agarose pucks were prepared by aspirating with a P10 pipette tip. Next, the pipette tip was lodged onto a stereological pipette tip allowing for a larger volume to be spun down onto the agarose puck. The fixed synaptosomes were spun onto the agarose puck using Labofuge 400R swinging bucket centrifuge at ~4'000g for 1 hour at 4 °C. Then, the liquid was aspirated carefully and the agarose puck was pushed out of the pipette tip and encapsulated in an agarose droplet. This droplet was stored at 4 °C in PBS until further processing.

Sorted synaptosomes were processed similarly to the above for EM with minor alterations. First, ~10 million synaptosomes were sorted into bovine albumin serum in PBS (final concentration 2%). The pipette tip containing the agarose puck was attached to a 15ml Falcon tube to accommodate the higher volume. Synaptosomes were spun down onto the agarose puck at 4 °C and 4000g for 3 hours after which glutaraldehyde was added to a final concentration of 2.5%. The synaptosomes were fixed at 4 °C at ~4000g for 30 minutes.

Next, samples were washed in 0,1 M Cacodylate buffer (CD) at room temperature and samples were further stained in 1% $OsO_4$ (in 0,1M CD) for 30 minutes. Samples were washed in 0,1M CD and water, and subsequently stained with 1% uranyl acetate (UA) (in water) in the dark for 10 minutes. Samples were washed with water and dehydrated by incubation in a series of ethanol buffers ranging from 30% to 100% ethanol.

For epoxy resin embedding, the samples were washed in dehydrated Propylenoxid and incubated in a 1:1 mixture of EPON:dehydrated Propylenoxid for 30 minutes at RT. Then, samples were left in EPON overnight at RT. The following day, excess agarose was trimmed and samples were placed in a silicone mold and EPON was polymerized for at least 48 hours at 60°C. 70 nm sections were generated using a Leica Leica Reichert Ultracut S Ultramicrotome with a DiATOME ultra 45° diamond knife and mounted onto a Pioloform film on copper grids. EM imaging was performed with a Zeiss LEO 912 AB Omega and a Sharp Eye TRS (2x2k) CCD camera. ImageSP was used to control the CCD camera and to make tilescans with 20% overlap at a magnification of 31'500X at 120 kEV for characterisation of F2/3 unsorted synaptosomes. FASS synaptosomes were imaged at 4100X magnification at 120 kEV. Tiles were stitched for visualization using TrakEM2 in ImageJ (Cardona et al. 2012). For image analysis, ~30 synaptosomes from each sample were randomly selected, blinded (using ImageJ plugin Blind

Analysis Tools), converted to 8-bit and the image contrast was normalized to 0.35 saturation. The blinded images were manually annotated for size, mitochondrial area and the presence of an attached postsynaptic density, while the number of SVs was determined with the automatic detection pipeline (Imbrosci, Schmitz, and Orlando 2022). Unblinding and subsequent analysis was performed in R. To estimate the true diameter, overcoming the random cross sectional nature of 2d TEM, the assumption of perfectly spherical synaptosomes was made. This simplification allows for sampling random cross sections from a sphere with a known radius (R) by solving for Y in $X^2 + Y^2 = R^2$ given randomly drawn X values from the range 0-R. Then 2Y can be used as the diameter of the randomly sampled cross section of the sphere. The above described was performed 30 times for four simulated samples.

<u>Reanalysis of scRNA data from Zeisel et. al.</u>

The datafile "l6_r2_cns_neurons.loom" (Zeisel et al. 2018) was analyzed using the R-package Seurat (Hao et al. 2021). Neurons were selected by dissection area and scGate (Andreatta, Berenstein, and Carmona 2022) was used to detect neurons expressing cre-driver line genes, Camk2a, Gad2 and Syn1, and excitatory/inhibitory markers vGlut1 and vGAT. Midbrain dopaminergic neurons were defined, using scGate, by Dat expression and combined by summing counts for neurons from the same mouse resulting in pseudobulk data. After normalizing for total counts, expression of genes of interest was plotted.

<u>Fluorescence-activated synaptosome sorting (FASS)</u>

FASS was performed as previously described (Biesemann et al. 2014; Hafner et al. 2019; Paget-Blanc et al. 2022). F2/3 fractions from the synaptosome preparation were diluted 1:50 in GM buffer and 1.5μg/ml membrane dye (FM4-64, Thermo Fisher) was added. Synaptosomes were analyzed and sorted on a FACSAria Fusion (BD Biosciences) running FACSDiva, equipped with a 70μm Nozzle and the following settings: 488nm laser (for FM4-64), 561nm laser (for TdTomato), sort precision (0-16-0), FSC (317 V), SSC (488/10 nm, 370V), PE "TdTomato" (586/15 nm, 470V), PerCP "FM4-64" (695/40 nm, 470), thresholds (FSC = 200, FM4-64 = 700). Samples were analyzed and sorted at approx. 20'000 events/s and a flow rate of < 3. Doublet particles were excluded based on SSC-H and SSC-W. We constructed a gate (P3) to obtain the synaptosome population that selects particles that are double-positive for TdTomato and FM4-64 by thresholding against synaptosomes from wt mice (Figure 1B). For control synaptosomes all non-doublet particles above the FM4-64 threshold were sorted (P2). For each sorted sample (P3) as well as the matching control sample (P2) we sorted 20 Mio particles (Experiment described in

Figure 1), 10 Mio particles (Experiment described in Figure 2A) or 2 Mio particles (Experiment described in Figure 6).

Synaptosome immunofluorescence

Sorted synaptosomes were centrifuged onto gelatin coated coverslips (Electron Microscopy Sciences) for 20 min at 4’000g and 4 °C in a Labofuge 400R and fixed for 15min in 4% PFA, 4% sucrose in PBS. Synaptosomes were then immunolabeled according to the manufacturer's instructions (Alexa Fluor 488 Tyramid SuperBoost Kit, Invitrogen, B40941) using the following antibodies: anti-PSD95 (Thermo Fisher, MA1-046, ms, 1:1000) and anti-Gephyrin (SYSY, 147011, ms, 1:1000), and subsequently mounted and imaged as described above for brain sections. Synaptosome co-localization analysis was performed in ImageJ using the SynaptosomesMacro published by Paget-Blanc et. al. (Paget-Blanc et al. 2022).

Synaptosome processing for MS

Sorted synaptosomes and control particles were filtered onto Whatman glass microfiber filters (GF/F, Cytiva) and stored at -80 °C until further processing. Samples from the experiment across brain regions were allocated to blocks containing one sample of each condition and batch-processed. Synaptosome samples with 10 Mio or 20 Mio particles were lysed in 20µl lysis buffer (100 mM Tris, 1% sodium deoxycholate, 10 mM TCEP, 15 mM 2-chloroacetamide) by repeated sonication using a VialTweeter (Hielscher Ultrasonics), and heated to 95 °C for 5 min. Proteins were digested with 0.1µg LysC and 0.1µg trypsin (Promega) overnight at 37 °C. Samples were then acidified with 10% formic acid to approximately pH 3, and centrifuged for 10 min at 16’000 g. The supernatant was desalted using ZipTip pipette tips (Merck) and dried in a vacuum centrifuge. Synaptosome samples with 2 Mio particles were solubilized in 20ul TEAB buffer (50mM Triethylammonium bicarbonate, 1mM $CaCl_2$, 0.05µg trypsin LysC and 0.05µg trypsin) and digested overnight at 37 °C. 30µl acetonitrile was added and the samples were centrifuged 10 min at 16’000 g. The supernatant was filtered through ZipTip pipette tips by centrifugation for 1 min at 2’000g and this step was repeated once after adding 50µl of 50% acetonitrile in MS-grade water to the pellet. Samples were dried in a vacuum centrifuge and stored at -20 until Liquid chromatography–tandem mass spectrometry (LC-MS/MS) analysis.

LC-MS/MS analysis

The peptide samples were reconstituted in 5% acetonitrile (ACN) and 0.1% formic acid (FA) supplemented with an iRT peptide standard (Biognosys). Peptide mixtures were analyzed using an UltiMate 3000 nano-LC coupled to a Fusion Lumos mass spectrometer (Thermo Fisher Scientific). Briefly, samples were loaded onto a PepMap 100 C18 trap column (75 µm id × 2 cm length, 3 µm particle size; Thermo Fisher Scientific) at 6 µL/min for 6 min with 2% acetonitrile (v/v) and 0.05% trifluoroacetic acid (v/v) followed by separation on an C18 analytical column (75 µm id × 50 cm length, 1.7 µm particle size; CoAnn Technologies) maintained at 55°C. Peptides were separated by a non-linear 120 min gradient (Muntel et al. 2019) using mobile phase A (100% H2O, 0.1% formic acid) and B (80% acetonitrile, 0.1% formic acid). The Fusion Lumos mass spectrometer was operated in DDA mode for spectral library generation and DIA mode for all samples used in this study, using acquisition methods that are described in Muntel et. al. (Muntel et al. 2019). In brief, the 40W DIA-method had the following settings: Full scan; orbitrap resolution = 120k, AGC target = 125%, mass range = 350-1650 m/z and maximum injection time = 100 ms. DIA scan; activation type: HCD, HCD collision energy = 27%, orbitrap resolution = 30k, AGC target = 2000%, maximum injection time = dynamic. The mass spectrometry proteomics data is currently being prepared for upload to the ProteomeXchange Consortium via the PRIDE (Perez-Riverol et al. 2019) partner repository and made available shortly.

Data analysis of DIA LC-MS/MS

LC-MS/MS DIA runs were analyzed with Spectronaut version 16 (Biognosys) as previously described (Bruderer et al. 2017; van Oostrum et al. 2020). Briefly, a spectral library was generated from collected in-house generated FASS synaptosome .raw files. The collected DDA spectra were searched against the UniProtKB/Swiss-Prot database for mus musculus using the Sequest HT search engine within Thermo Proteome Discoverer 2.4 (Thermo Fisher Scientific). The identified proteins were assessed using Percolator and filtered using the high peptide confidence setting in Proteome Discoverer. Analysis results were then imported to Spectronaut for the generation of spectral libraries.

Targeted data extraction of DIA-MS acquisitions was performed in Spectronaut with default settings. The proteotypicity filter “only protein group specific” was applied. Extracted features were exported from Spectronaut for statistical analysis with MSstats 4 (Choi et al. 2014) using default settings. Briefly, for each protein, features were log-transformed and fitted to a mixed effect linear regression model for each sample. The model estimated fold change and statistical significance for all compared conditions. For the P2 control conditions, all P2 samples were grouped according to the brain region of origin independent of the mouse line. Benjamini–Hochberg method was used to account for multiple testing and p-value adjustment was performed on all proteins that met the fold-change cutoff. Significantly different proteins were determined using the threshold log2 fold-change >1.1 and adjusted p-value < 0.05. For analysis comparing conditions originating from the same brain region the significance testing result and sample quantification (protein abundance) files from MSstats were used. For analysis comparing multiple conditions across brain regions we calculated the normalized log2(fold-change) of the matched P3/P2 pairs for every sample and protein, which was then used for further analysis.

Data analysis of synaptic proteomes

The synaptic proteomes for Camk2a+, Gad2+ and Syn+ proteomes were determined by quantitative enrichment against the P2 control samples and the other synapse types originating from the same brain region. Specifically, the proteomes were defined as proteins that meet the fold-change and p-value cutoff comparing P3 with P2 samples of the same brain region. Proteins that meet the cutoff (P3/P2) in several of the cre-driver lines within the same brain region were allocated to all synapses types were they met the cutoff unless they were significantly enriched in one over the other in the direct comparison, in that case the protein was assigned only to the cre-driver line where it was found enriched. Proteins that were significantly enriched in one synapse type compared to another of the same brain region, and additionally had a positive fold-change compared to the control samples, were also included in the synaptic proteome of the first synapse type. The Dat+ and cortical interneuron proteomes were defined by quantitative enrichment against the P2 control samples using the above mentioned cutoffs. All further analysis was done based on quantitative values obtained from MSstats in the R environment. Variance partitioning was analyzed using the VariancePartition R package (Hoffman and Schadt 2016). The chord diagram was generated using the circlize R package (Gu et al. 2014). Protein to gene name conversion was done using either org.Mm.eg.db (Carlson, n.d.) or the UniProt API. SynGO analyses were performed using the SynGO web page with default settings (Koopmans et al. 2019). Euler diagrams were constructed using the eulerr package (Larsson and Gustafsson 2018). STRING (Szklarczyk et al. 2019) networks were generated in Cytoscape using the Cytoscape StringApp (Doncheva et al. 2019) and analyzed using CentiScaPe (Scardoni, Petterlini, and Laudanna 2009). Protein complexes were annotated using the CORUM database (Giurgiu et al. 2019). Weighted gene correlation network analysis was done using the WGCNA R package (Langfelder and Horvath 2008) using soft power 6, a signed hybrid network type and the bicor correlation function. Heatmaps were visualized using ComplexHeatmap (Gu, Eils, and Schlesner 2016). The adjacency matrix was visualized as protein-protein correlation network in Cytoscape with a frequency cutoff of 0.3. Gene set enrichment analysis (GSEA) for Gene Ontology terms (“The Gene Ontology Resource: Enriching a GOld Mine” 2021) and KEGG pathways (Kanehisa et al. 2023) was performed with clusterprofiler (Wu et al. 2021). ggplot2 was used for visualization (Wickham 2016). Cartoons were generated using Adobe Illustrator or Biorender.com. FACS plots were generated with FlowJo.

**Method references**


Andreatta, Massimo, Ariel J. Berenstein, and Santiago J. Carmona. 2022. "scGate: Marker-Based Purification of Cell Types from Heterogeneous Single-Cell RNA-Seq Datasets." *Bioinformatics* 38 (9): 2642–44.

Bäckman, Cristina M., Nasir Malik, Yajun Zhang, Lufei Shan, Alex Grinberg, Barry J. Hoffer, Heiner Westphal, and Andreas C. Tomac. 2006. "Characterization of a Mouse Strain Expressing Cre Recombinase from the 3' Untranslated Region of the Dopamine Transporter Locus." *Genesis* 44 (8): 383–90.

Biesemann, Christoph, Mads Grønborg, Elisa Luquet, Sven P. Wichert, Véronique Bernard, Simon R. Bungers, Ben Cooper, et al. 2014. "Proteomic Screening of Glutamatergic Mouse Brain Synaptosomes Isolated by Fluorescence Activated Sorting." *The EMBO Journal* 33 (2): 157–70.

Bruderer, Roland, Oliver M. Bernhardt, Tejas Gandhi, Yue Xuan, Julia Sondermann, Manuela Schmidt, David Gomez-Varela, and Lukas Reiter. 2017. "Optimization of Experimental Parameters in Data-Independent Mass Spectrometry Significantly Increases Depth and Reproducibility of Results." *Molecular & Cellular Proteomics: MCP*, October. https://doi.org/10.1074/mcp.RA117.000314.

Cardona, Albert, Stephan Saalfeld, Johannes Schindelin, Ignacio Arganda-Carreras, Stephan Preibisch, Mark Longair, Pavel Tomancak, Volker Hartenstein, and Rodney J. Douglas. 2012. "TrakEM2 Software for Neural Circuit Reconstruction." *PloS One* 7 (6): e38011.

Carlson, M. n.d. "Org. Mm. Eg. Db: Genome Wide Annotation for Mouse. R Package Version 3.2. 3." *. London, United Kingdom: Genome Biology (BMC)*.

Choi, Meena, Ching-Yun Chang, Timothy Clough, Daniel Broudy, Trevor Killeen, Brendan MacLean, and Olga Vitek. 2014. "MSstats: An R Package for Statistical Analysis of Quantitative Mass Spectrometry-Based Proteomic Experiments." *Bioinformatics* 30 (17): 2524–26.

Doncheva, Nadezhda T., John H. Morris, Jan Gorodkin, and Lars J. Jensen. 2019. "Cytoscape StringApp: Network Analysis and Visualization of Proteomics Data." *Journal of Proteome Research* 18 (2): 623–32.

Giurgiu, Madalina, Julian Reinhard, Barbara Brauner, Irmtraud Dunger-Kaltenbach, Gisela Fobo, Goar Frishman, Corinna Montrone, and Andreas Ruepp. 2019. "CORUM: The Comprehensive Resource of Mammalian Protein complexes—2019." *Nucleic Acids Research*. https://doi.org/10.1093/nar/gky973.

Gu, Zuguang, Roland Eils, and Matthias Schlesner. 2016. "Complex Heatmaps Reveal Patterns and Correlations in Multidimensional Genomic Data." *Bioinformatics* 32 (18): 2847–49.

Gu, Zuguang, Lei Gu, Roland Eils, Matthias Schlesner, and Benedikt Brors. 2014. "Circlize Implements and Enhances Circular Visualization in R." *Bioinformatics* 30 (19): 2811–12.

Hafner, Anne-Sophie, Paul G. Donlin-Asp, Beulah Leitch, Etienne Herzog, and Erin M. Schuman. 2019. "Local Protein Synthesis Is a Ubiquitous Feature of Neuronal Pre- and Postsynaptic Compartments." *Science* 364 (6441). https://doi.org/10.1126/science.aau3644.

Hao, Yuhan, Stephanie Hao, Erica Andersen-Nissen, William M. Mauck 3rd, Shiwei Zheng,

Andrew Butler, Maddie J. Lee, et al. 2021. “Integrated Analysis of Multimodal Single-Cell Data.” *Cell* 184 (13): 3573–87.e29.
Hippenmeyer, Simon, Eline Vrieseling, Markus Sigrist, Thomas Portmann, Celia Laengle, David R. Ladle, and Silvia Arber. 2005. “A Developmental Switch in the Response of DRG Neurons to ETS Transcription Factor Signaling.” *PLoS Biology* 3 (5): e159.
Hoffman, Gabriel E., and Eric E. Schadt. 2016. “variancePartition: Interpreting Drivers of Variation in Complex Gene Expression Studies.” *BMC Bioinformatics* 17 (1): 483.
Imbrosci, Barbara, Dietmar Schmitz, and Marta Orlando. 2022. “Automated Detection and Localization of Synaptic Vesicles in Electron Microscopy Images.” *eNeuro* 9 (1). https://doi.org/10.1523/ENEURO.0400-20.2021.
Jong, Annemieke de, Karianne G. Schuurman, Boris Rodenko, Huib Ovaa, and Celia R. Berkers. 2012. “Fluorescence-Based Proteasome Activity Profiling.” *Methods in Molecular Biology* 803: 183–204.
Kanehisa, Minoru, Miho Furumichi, Yoko Sato, Masayuki Kawashima, and Mari Ishiguro-Watanabe. 2023. “KEGG for Taxonomy-Based Analysis of Pathways and Genomes.” *Nucleic Acids Research* 51 (D1): D587–92.
Koopmans, Frank, Pim van Nierop, Maria Andres-Alonso, Andrea Byrnes, Tony Cijsouw, Marcelo P. Coba, L. Niels Cornelisse, et al. 2019. “SynGO: An Evidence-Based, Expert-Curated Knowledge Base for the Synapse.” *Neuron* 103 (2): 217–34.e4.
Langfelder, Peter, and Steve Horvath. 2008. “WGCNA: An R Package for Weighted Correlation Network Analysis.” *BMC Bioinformatics* 9 (December): 559.
Larsson, Johan, and Peter Gustafsson. 2018. “A Case Study in Fitting Area-Proportional Euler Diagrams with Ellipses Using Eulerr.” In *SetVR@ Diagrams*, 84–91.
Muntel, Jan, Joanna Kirkpatrick, Roland Bruderer, Ting Huang, Olga Vitek, Alessandro Ori, and Lukas Reiter. 2019. “Comparison of Protein Quantification in a Complex Background by DIA and TMT Workflows with Fixed Instrument Time.” *Journal of Proteome Research*, February. https://doi.org/10.1021/acs.jproteome.8b00898.
Oostrum, Marc van, Benjamin Campbell, Charlotte Seng, Maik Müller, Susanne Tom Dieck, Jacqueline Hammer, Patrick G. A. Pedrioli, Csaba Földy, Shiva K. Tyagarajan, and Bernd Wollscheid. 2020. “Surfaceome Dynamics Reveal Proteostasis-Independent Reorganization of Neuronal Surface Proteins during Development and Synaptic Plasticity.” *Nature Communications* 11 (1): 4990.
Paget-Blanc, Vincent, Marlene E. Pfeffer, Marie Pronot, Paul Lapios, Maria-Florencia Angelo, Roman Walle, Fabrice P. Cordelières, et al. 2022. “A Synaptomic Analysis Reveals Dopamine Hub Synapses in the Mouse Striatum.” *Nature Communications* 13 (1): 3102.
Perez-Riverol, Yasset, Attila Csordas, Jingwen Bai, Manuel Bernal-Llinares, Suresh Hewapathirana, Deepti J. Kundu, Avinash Inuganti, et al. 2019. “The PRIDE Database and Related Tools and Resources in 2019: Improving Support for Quantification Data.” *Nucleic Acids Research* 47 (D1): D442–50.
Scardoni, Giovanni, Michele Petterlini, and Carlo Laudanna. 2009. “Analyzing Biological Network Parameters with CentiScaPe.” *Bioinformatics* 25 (21): 2857–59.
Sebring, R. J., N. F. Johnson, and W. D. Spall. 1988. “Transmission Electron Microscopy of Small Numbers of Sorted Cells.” *Cytometry* 9 (1): 88–92.
Szklarczyk, Damian, Annika L. Gable, David Lyon, Alexander Junge, Stefan Wyder, Jaime Huerta-Cepas, Milan Simonovic, et al. 2019. “STRING v11: Protein-Protein Association Networks with Increased Coverage, Supporting Functional Discovery in Genome-Wide Experimental Datasets.” *Nucleic Acids Research* 47 (D1): D607–13.
Taniguchi, Hiroki, Miao He, Priscilla Wu, Sangyong Kim, Raehum Paik, Ken Sugino, Duda Kvitsiani, et al. 2011. “A Resource of Cre Driver Lines for Genetic Targeting of GABAergic Neurons in Cerebral Cortex.” *Neuron* 71 (6): 995–1013.
“The Gene Ontology Resource: Enriching a GOld Mine.” 2021. *Nucleic Acids Research* 49 (D1):

D325–34.
Tsien, Joe Z., Dong Feng Chen, David Gerber, Cindy Tom, Eric H. Mercer, David J. Anderson, Mark Mayford, Eric R. Kandel, and Susumu Tonegawa. 1996. "Subregion- and Cell Type–Restricted Gene Knockout in Mouse Brain." *Cell* 87 (7): 1317–26.
Westmark, Pamela R., Cara J. Westmark, Athavi Jeevananthan, and James S. Malter. 2011. "Preparation of Synaptoneurosomes from Mouse Cortex Using a Discontinuous Percoll-Sucrose Density Gradient." *Journal of Visualized Experiments: JoVE*, no. 55 (January): e3196–e3196.
Wickham, Hadley. 2016. "ggplot2: Elegant Graphics for Data Analysis Springer-Verlag New York; 2009." *Preprint at*.
Wu, Tianzhi, Erqiang Hu, Shuangbin Xu, Meijun Chen, Pingfan Guo, Zehan Dai, Tingze Feng, et al. 2021. "clusterProfiler 4.0: A Universal Enrichment Tool for Interpreting Omics Data." *Innovation (Cambridge (Mass.))* 2 (3): 100141.
Zeisel, Amit, Hannah Hochgerner, Peter Lönnerberg, Anna Johnsson, Fatima Memic, Job van der Zwan, Martin Häring, et al. 2018. "Molecular Architecture of the Mouse Nervous System." *Cell* 174 (4): 999–1014.e22.
Zhu, Y., M. I. Romero, P. Ghosh, Z. Ye, P. Charnay, E. J. Rushing, J. D. Marth, and L. F. Parada. 2001. "Ablation of NF1 Function in Neurons Induces Abnormal Development of Cerebral Cortex and Reactive Gliosis in the Brain." *Genes & Development* 15 (7): 859–76.
